# Small patches dominate the Natura 2000 network of protected areas in Europe

**DOI:** 10.64898/2026.06.04.730078

**Authors:** Jutta Beher, Piero Visconti

## Abstract

The Natura 2000 network for protected areas is one of Europe’s main tool to address the loss of biodiversity. Voluntary commitments of all 27 Member States aim for an extension towards a coherent network of protected land and sea area by 2030 that covers at least 30% of each realm in an ecologically representative and spatially coherent way. Spatial coherence refers to addressing issues brought about by small size of protected patches and their fragmentation in the Natura 2000 network, which, however have not been analysed in depth. This study presents an analysis of the full Natura 2000 data set of terrestrial and marine protected areas in Europe, to which extent documented sites are split up into smaller fragments, how their spatial configuration and shape might mitigate or invite edge effects, and which edge effects might be expected based on the surrounding land cover and land use matrix. Our results show a substantial fragmentation of the network, with four times more individual patches than recorded sites, and 75% of these patches being smaller than 1 km^2^. Across four size categories between smaller than 1 km^2^ and larger than 100 km^2^, about a quarter of patches show a spatial configuration that is likely to cause substantial edge effects due to a suboptimal ratio between area and perimeter. At the same time, many of the smallest patches are surrounded by a large share of sealed surfaces and agricultural areas within a 500m radius. Considering fragmentation and edge effects in future reporting, monitoring and planning of additional designations and restoration activities could close an important gap in understanding causes for continuing declines of Europe’s biodiversity and highlight pathways to mitigate them.

**Graphical Abstract:** 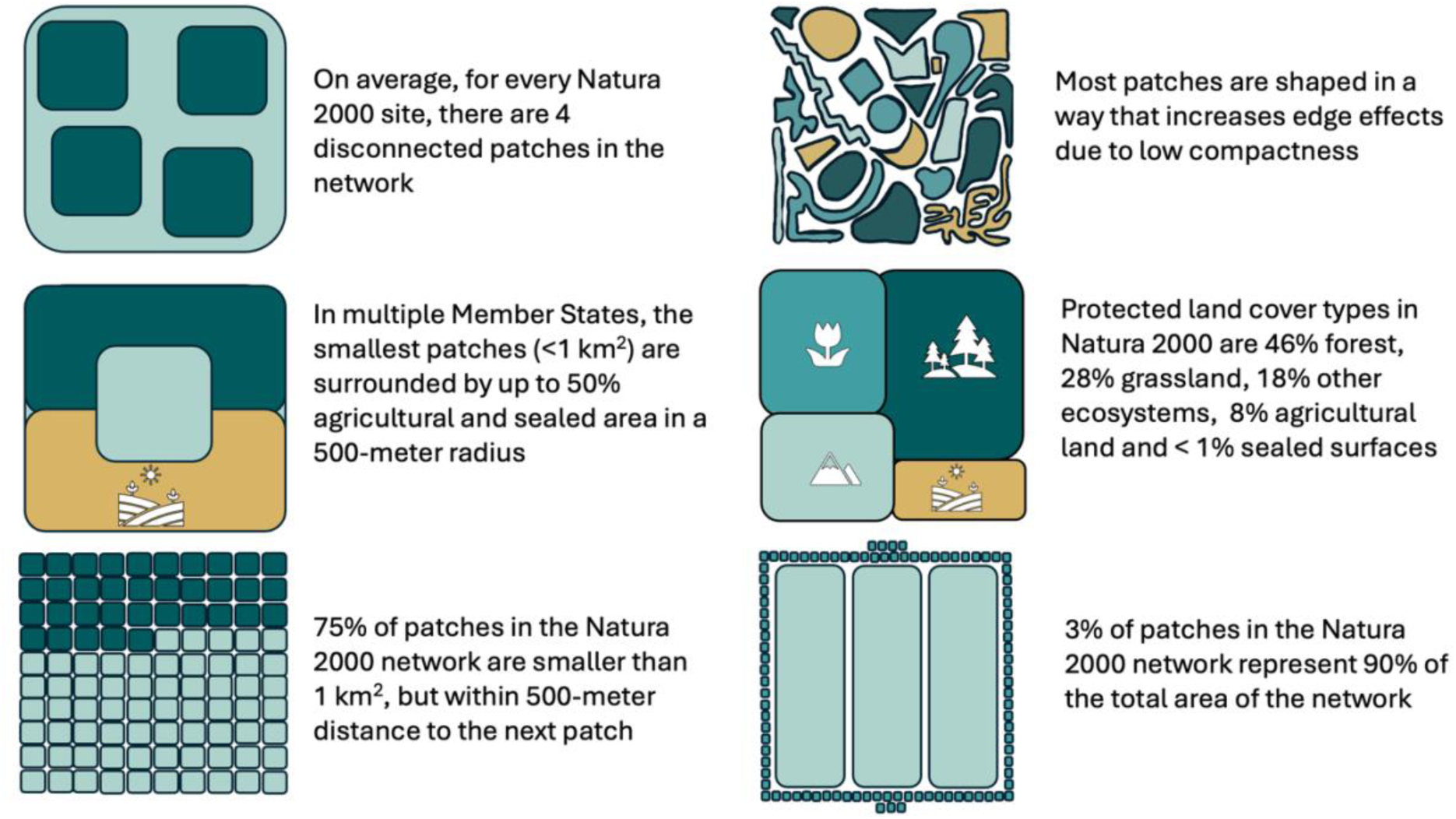

## Introduction

### Background

The establishment and management of protected areas are some of the most implemented conservation actions around the world (Maxwell et al., 2020). Europe is no exception, having established targets related to protected area designation and management as the first three targets of the EU Biodiversity Strategy to 2030 (European Commission, 2020). These include expanding the protected area network towards a coherent and resilient Trans-European Nature Network that is ecologically representative and cover at least 30% of the terrestrial and marine realm (Target 1); strictly protect at least 10% of marine and terrestrial extent of the EU (Target 2); and effectively manage all protected areas (Target 3). These targets build on two long-standing legal instruments that protect wild species and natural habitats are legally protected via directive 2009/147/EC (known as “Birds Directive”) and directive 92/43/EEC (known as the “Habitats Directive”), with several voluntary commitments of Member states, as requested by in the Biodiversity Strategy 2030 (European Commission, 2020; European Commission: Directorate-General for Environment and Sundseth, K, 2025). These two, so-called ‘’Nature Directives’’, also require EU Member states to designate and manage a network of international protected areas: Natura 2000 sites, to protect habitat and species in particular need of area-based conservation measures, listed in Annex I and II of the Habitats Directive and Annex II of the Birds Directive, in proportion to the representation within their territory, to maintain or, where appropriate, restore the conservation status in their natural range to favorable (European Economic Community (EEC), 1992), article 3).

However, despite increasing protected area coverage, the most recent reporting period shows species populations are mostly declining or stable, including inside Protected Areas and despite the relative benefit of EU protected areas, as the difference in population trends against comparable unprotected sites is statistically undetectable (EEA, 2020; European Commission. Joint Research Centre. & European Environment Agency., 2025; European Environment Agency., 2025; Gregory et al., 2019). The negative trends for many species and habitats are expected to continue (European Environment Agency., 2025).

Many factors concur to the lack of effectiveness of Protected Areas in mitigating threats to biodiversity. The key problems are in general well described for each taxa and habitat, including persistent pressures of land use and land use change, pollution, climate change, alien invasive species and overexploitation (European Environment Agency., 2025). While broad concepts and objectives are necessary and essential, the success and failure of any measures taken, including the designation of protected areas and management of existing pressures within and around them, will depend on the local context. For example, spatial characteristics of protected areas such as their size, isolation and fragmentation, as well as surrounding land use, can have a strong impact on intensity and presence of threats to species and habitats. Direct implications include edge effects, such as increased mortality or other issues caused by edge-related pollution, invasive or alien species, human disturbance and altered environmental conditions that impact protected areas but originate in the surrounding landscape “matrix” (Fletcher et al., 2024; Porensky & Young, 2013; Woodroffe & Ginsberg, 1998). Indirect implications include fragmentation and isolation causing loss of functional connectivity, which can limit genetic exchange and movement of species which in turn impact reproduction rates and adaptive potential of populations (Crooks et al., 2017; Fletcher et al., 2024; Gutiérrez-Rodríguez et al., 2017; Hanson et al., 2019, 2022).

However, in addition to the lack of available funding to implement conservation measures, the large gaps of detailed monitoring data as well as a large number of missing management plans at the site level means that site specific knowledge for each protected area does not necessarily exist to guide the most impactful management interventions to mitigate threats effectively on the local level (European Commission. Joint Research Centre. & European Environment Agency., 2025; European Environment Agency., 2020; Hermoso et al., 2022; Moersberger et al., 2024).

Few studies have investigated the level of fragmentation of the EU protected area network (EEA, 2022; Kubacka et al., 2022; Lawrence et al., 2021; Lawrence & Beierkuhnlein, 2023). Results based on data from 2011 and using a resolution of 1km^2^, found that habitat types show high levels of fragmentation in their spatial configuration in and around Natura 2000 sites across all of Europe, with a quarter of protected areas classified as highly fragmented (Lawrence et al., 2021). However, sites smaller than 1km^2^ were excluded, and the analysis did not contain aggregated results at the Member State level. To date, neither the size distribution of Natura 2000 sites nor the distances between them have been systematically quantified across the network. This represents an important knowledge gap, particularly because many sites consist of multiple small patches embedded within landscapes comprising diverse land-use types. Despite evidence in the last State of Nature report suggesting that many protected areas may fail to achieve conservation objectives due to edge effects and the acknowledgment that the establishment and management of buffer zones around protected areas is a global and local challenge (European Environment Agency, 2020, quote page 124: "*Furthermore, although site-level management is sufficient in many countries, conservation objectives are sometimes not being met due to pressures from outside the sites (e.g. nitrogen pollution or changes in hydrological regimes). The effective implementation of other directives is therefore critical, as well as an increased policy focus on establishing effective buffer zones and semi-natural habitats outside protected areas and ensuring adequate levels of protection inside sites*"), the extent to which EU protected areas include and are surrounded by human-dominated environment has also not been systematically analysed and synthesised.

To address these knowledge gaps, we present data for the European Union summarising information on patch size, compactness, and fragmentation and isolation of Natura 2000 sites as well as land-use within and around them. We also disaggregate our results for EU Member States, describing different patterns emerging for each indicator across countries. We conclude the implications of these findings for species conservation and possible improvements in design and management of protected areas.

## Methods

### Extent of the analyses and data sources

The analyses cover all 27 countries of the European Union, using data of administrative boundaries, protected areas, and land use (Table S1). To delimit administrative boundaries, we used data from Eurostat (Eurostat, 2024). The spatial boundaries of Natura 2000 sites were used from the year 2024, the most up-to-date version of the data at the time of analysis, including areas documented up to 2022 (European Environment Agency & European Commission, 2025). Spatial analyses including other national designations of protected areas used the “Nationally Designated Areas (NatDa) dataset (formerly CDDA, version of 2023 (European Environment Agency, 2024)). Satellite data on land use and landcover was used at a 10 meter resolution based on the year 2023 (Copernicus Land Monitoring Service & Copernicus Land Monitoring Service Helpdesk, 2025). All analyses were realised in python and R, using the arcpy package and sf, tidyverse, GridExtra and stringr packages. Figures were produced with python (arcpy, plotly and pandas) and R (tidyverse), maps in ArcPRO. All code is publicly available on figshare (private link: https://figshare.com/s/05fd3fcf8fd763735f82)

### Spatial analysis workflow

Three separate analyses were performed: 1) quantification of the extent of size distribution and isolation of Natura 2000 sites, 2) quantification of land use and land cover within and around Natura 2000 sites, and 3) a worked example of how to inform assessment and planning for improved conservation management outcomes on the species or habitat level (Figure 1).

**Figure 1.**
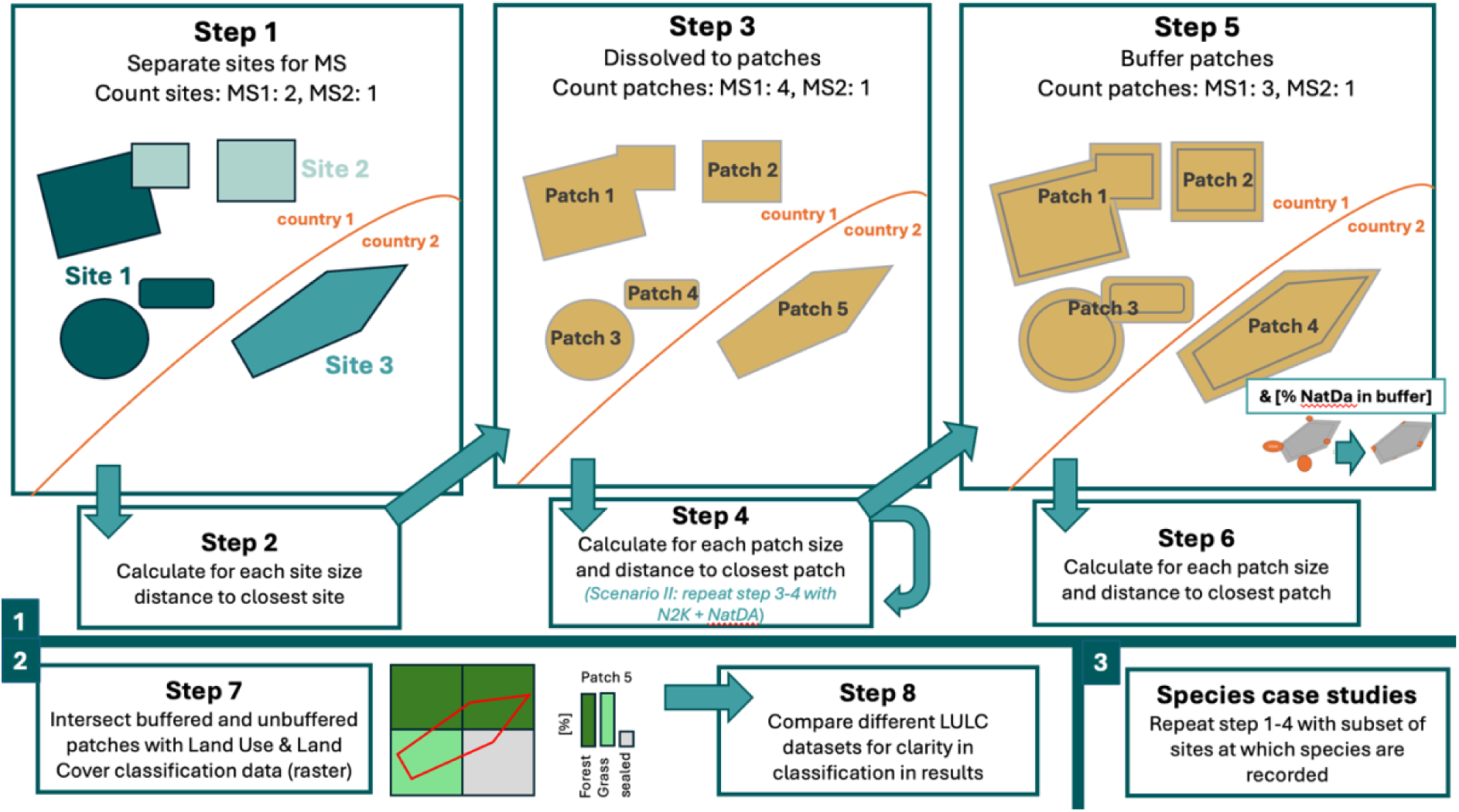
workflow of individual steps for three aspects of the fragmentation analysis. In the first aspect, protected area sites were separated for each EU Member State, and number, individual site area extent, and distance to the next closest site were calculated. Then, sites were aggregated into individual polygons (patches), and the calculations were repeated. Then, individual patches were surrounded with a 500-meter-wide buffer, and all calculations were repeated with the enlarged and merged patches. In the second aspect, land use categories within individual patches and the immediate surroundings in the buffer zone outside of the protected areas were extracted from available satellite data. To illustrate the relevance for conservation management assessment and planning, as a third aspect, the analysis was applied to three example species and the findings related to their ecology and conservation status.

#### Natura 2000 sites size distribution, shape, and isolation of individual patches

To enable a time efficient analysis workflow, Natura 2000 sites were extracted as individual layers for each Member State using the individual two-letter country codes. In a first run of the workflow, we limited the data to terrestrial protected areas by clipping Natura 2000 sites with administrative boundaries, but this resulted in many artifacts due to mismatches between delineations of outlines, including marine protected areas and coastlines. To avoid the artificial inflation of small patches during the clipping process, we then ran the workflow with the full dataset including terrestrial and marine sites.

In the first step of the workflow, we calculated the total extent of each Natura 2000 site in hectares, as well as the distance to the closest next site. In the second step, we separated the individual (multi-part) polygon of each site into individual (single-part) patches to quantify existing fragmentation of the documented sites. The third step filtered sites and patches to drop any polygon smaller than 5 m^2^ to account for the involuntary generation of small slivers due to crossing outlines as an artefact of the spatial analysis process, and other areas where the size of the polygon does not reflect the size of used habitat (such as entrances to caves). In the last step, we calculated for each Member State the total number of the patches produced, as well as their area in hectare, a metric for compactness (roundness = perimeter^2 / (4 * pi * area), and distance to closest next (single feature) polygon patch.

While several national designations are not always managed for biodiversity conservation objectives, e.g. Landschaftsschutzgebiete in Germany, they could potentially mitigate against direct threats to biodiversity, such as human disturbance, intensive land-use, and unsustainable hunting, and act as connecting areas between Natura 2000 sites. Therefore, we repeated the four-step analysis by considering both, using merged data of the Natura 2000 network and Nationally Designated Protected Areas (NatDa) in a second scenario.

To allow insights into potential issues of land cover and use in the immediate surroundings of protected areas and required spatial scales for mitigating fragmentation, a buffer distance was selected that would bridge the gaps between the majority of individual patches. A calculation of all distances to the closest next patch showed that a buffer size of 500 meters captures about 75% of all distances between each of the smallest patches to their closest next patch (Figure S1). We applied a buffer of 500-meter radius to each patch and repeated the four-step analysis, calculating the total number, as well as the size of each patch and distance to the closest next patch. Additionally, we calculated the percentage of Nationally Designated Protected Areas (NatDa dataset) within the buffer zone to assess the function of national designations as a protective and connecting layer in the Natura 2000 network. All analyses at the country level are based on the documented country code in the Natura 2000 data set.

#### Land use and land cover in and around Natura 2000 area

To derive information on the land cover and use within and around Natura 2000 sites, classification of land use cover was extracted from the CLCplus Backbone 10m raster data for all individual patches and quantified in hectares for each Member State. The data was visualised as a treemap, both for the area within Natura 2000 patches, and the area in the buffer around the Natura 2000 patches. CLCplus at 10-meter resolution has 12 classes (Table S6). In contrast to the first part of the analysis, land cover categories were extracted for terrestrial and freshwater patches, because the satellite data does not cover offshore areas and would, therefore, only cover a fraction of the marine protected areas. The category of ocean was therefore eliminated from the data before plotting Satellite data has known limitations in detecting and classifying the intensity of human land use, particularly in grasslands, and classification might also not match the exact habitat categories used in European conservation management. Because of these limitations, and the importance of distinguishing natural from anthropogenic land cover to assess impacts of fragmentation, we tested the sensitivity of our findings by comparing the 10m CLCplus Backbone raster data with the Copernicus Land Monitoring Service (CLMS) N2K 2018 vector product (Table S1, Supplementary Material S6), for which land cover has been validated for 8 land cover categories and 55 subcategories, for a subset of grassland rich Natura 2000 sites across Europe. We applied the comparison in one country, Austria, for all patches existing in both datasets (Supplementary Material S5: Accuracy of land use data). Based on this sensitivity analysis, the 12 classes of the CLCplus Backbone imagery were summarised into 5 broader but more robust categories. Four categories showed a good match with the more accurate vector layer, and one category was more difficult to match:

1. forest types (evergreen, needle, broadleaved), high accuracy
2. grassland (permanent herbaceous), lower accuracy likely due to difficulty in distinguishing managed and unmanaged grasslands and some agricultural fields
3. other non-forested natural areas (low growing woody vegetation, sparse/non vegetated, water, snow/ice), high accuracy
4. sealed surfaces, high accuracy
5. periodically herbaceous (= likely agriculture), high accuracy

#### Application of the analysis for species and habitat-specific conservation assessment and planning

To provide an assessment of existing risk of edge effects and fragmentation at the species level, and a first assessment of the potential underestimation of fragmentation in the general analysis, we chose three species that depend on grasslands, including agricultural areas and pastures. Little bustard *Tetrax terax* and meadow viper *Vipera ursinii* are listed as vulnerable in the IUCN redlist, while Pannonian tundra vole *Microtus oeconomus mehely* is one of the most endangered small mammals in Europe. All Natura 2000 site ID codes for which the species is reported were extracted from the Natura 2000 viewer. Based on the ID codes indicating species’ presence, we subset the spatial data set of Natura 2000 sites, and quantified patches following the described methods above (Step 9 in Figure 1), complemented by a visual inspection of several locations in QGIS. Because it is not possible to know if a species or habitat can be found at only one, several or all patches of any given site because species records are not spatially explicit at a finer scale, we did not include the extraction of land cover here, as results would be impossible to interpret.

## Results

### Natura 2000 sites’ size distribution and isolation of individual patches

The Natura 2000 dataset includes a total of 27.020 individual sites (26.679 individual sites when clipped to the terrestrial area of Europe to exclude marine sites), which translate into 55.460 individual patches (Figure 2a, Figure 2b). The number of patches increased to 107.632 when that Natura 2000 dataset was clipped to the terrestrial area of Europe, due to mismatching delineations of coastline and marine protected area boundaries, creating small cut-off polygons from the marine protected area on land (Figure S2). In the remainder of the manuscript, we describe results for the analysis of the size of patches of the full Natura 2000 dataset, without clipping, but focus on terrestrial land cover and land use only. At the site level, the smallest size category up to 1km^2^ includes 9493 patches (∼37%), followed by 8821 sites between 1 and 10 km^2^ (∼32%), 6120 sites between 10 and 100 km^2^ (∼22%) and 2586 sites larger than 100 km^2^ (∼7%) (Figure 2a). At the patch level, the relative frequency of the different sizes changes significantly, percentagewise, there are double the amount of the smallest size category compared to sites (40.840 patches, ∼74%), half the size category between 1 and 10 km^2^ (9945 patches, ∼18%), a quarter of the category between 10 and 100 km^2^ (3490 patches, ∼6%) and less than a third of the largest category (1185 patches, ∼2%) (Figure 2b). Expanding each patch by 500m decreased the number of individual disconnected patches from 55.460 to 18.154, less than the total number of reported sites, as a result of overlapping buffered patches being merged together (Figure 2: Number and size of sites (a) and patches (b) of the Natura 2000 network in the European Union. Percent is shown along the y-axis, and count of patches and sites is shown as labels on each bar. Analysis at the site level underestimates the number small-sized class of land under protection that are surrounded by other land management types by a factor of 4 and overestimates the largest size group by a factor of 2.Figure 2c). This means that 98% of protected area patches are within 1km distance of the next closest patch (see also Figure S1). Considering Nationally Designated Protected Areas in the analysis of individual patches of the Natura 2000 network increases the percent and count of patches smaller than 1km^2^ substantially to 90% at the EU level, adding over 200.000 additional patches in the smallest size category, while the other size classes stay relatively similar considering the count of patches, but greatly reduce in proportion to the total (Figure 2d). The percentage of national designations within the 500-meter buffer area around Natura 2000 sites is 12.3% on average across all EU Member States.

**Figure 2.**
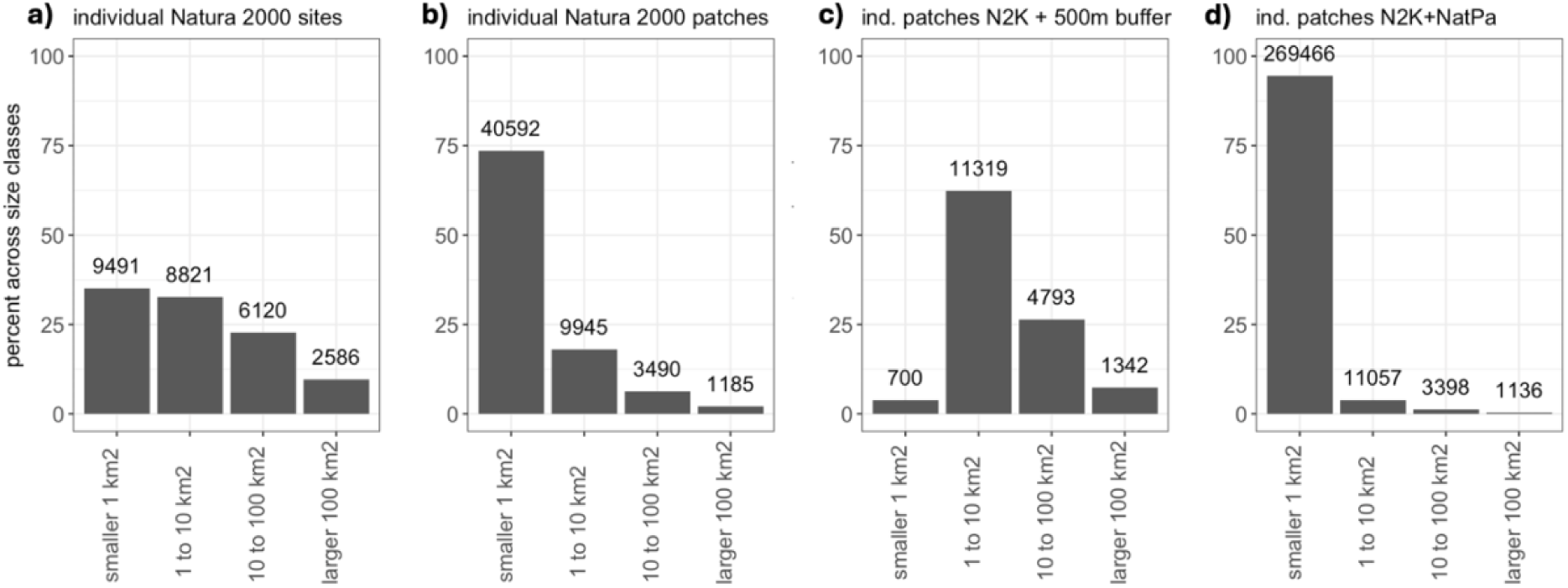
Number and size of sites (a) and patches (b) of the Natura 2000 network in the European Union. Percent is shown along the y-axis, and count of patches and sites is shown as labels on each bar. Analysis at the site level underestimates the number small-sized class of land under protection that are surrounded by other land management types by a factor of 4 and overestimates the largest size group by a factor of 2. Buffering patches with 500m (c) reduces the amount of patches smaller than 1km significantly, while accounting for Nationally Designated Protected Areas (d) increases the smallest size category significantly in percent and number.

The size distribution of sites less than 1 km^2^ across 5 size categories from 5 m^2^ to 1 km^2^ shows only 50 sites smaller than 100 m^2^, with an exponential increase across the larger categories and over 50% larger than 0.1 km^2^ (100.000m^2^). At the patch level, there are also least patches in the smallest categories and most in the largest, but the increase is less pronounced, with around 35% in the largest group, and more than 1000 patches smaller than 100 m^2^, a 24-fold increase compared to the site level (Figure 3).

**Figure 3.**
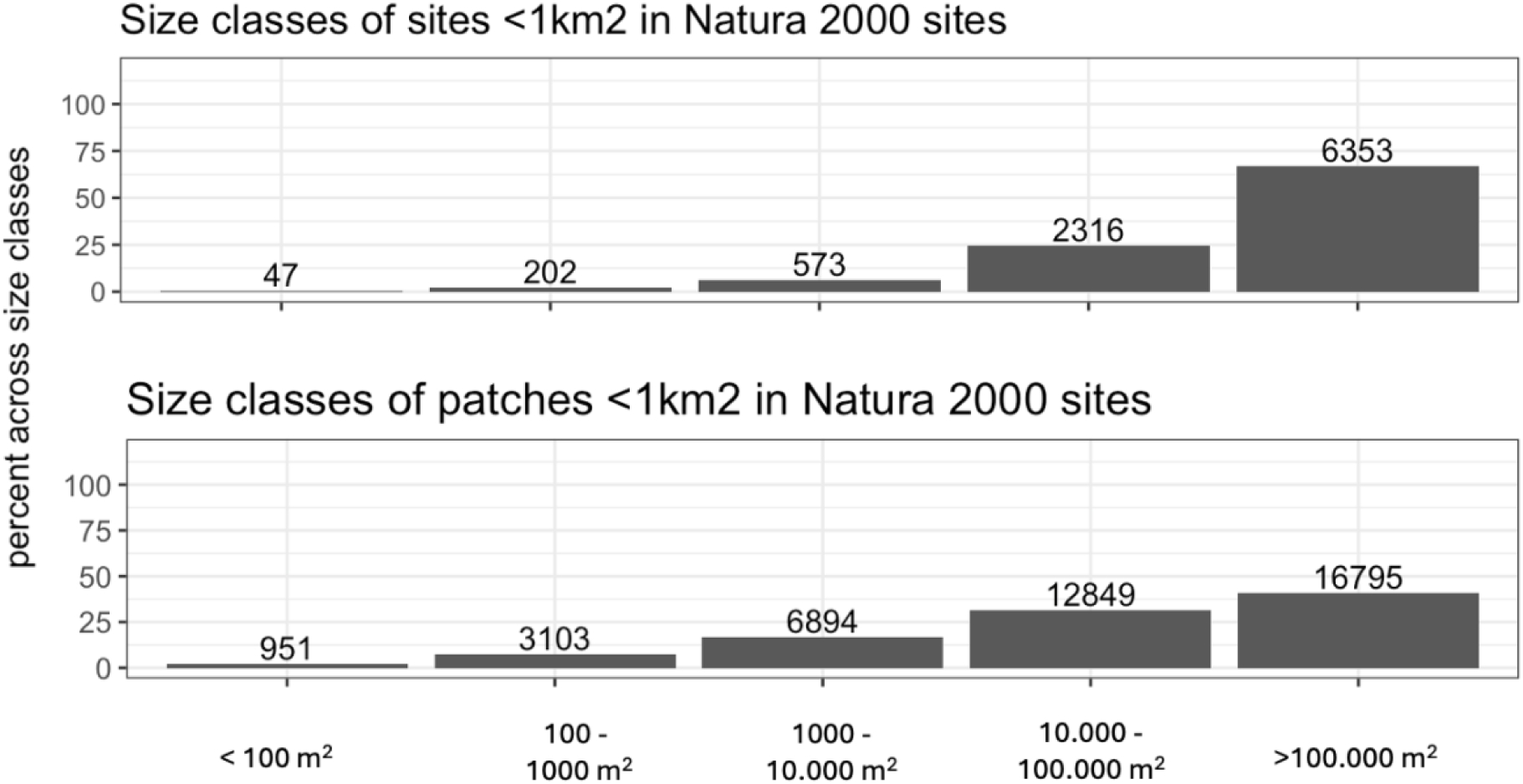
Size distribution of protected area sites (top) and patches (bottom) as percent of the total count of patches along the y-axis. All bars are labelled with the count of patches in each size category.

When disaggregating our results at country level, we find that the relative higher frequency of sites smaller than 1km^2^ are particularly prevalent in Austria, Czech Republic, Croatia, Malta, Sweden, Slovenia, and Slovakia, where half or more of all sites fall in this category. Germany and Sweden stand out with more than a thousand sites smaller than 1km^2^ (Figure 4 and Figure S4a). At the patch level, the smallest size category (<1km^2^) is most common in all countries except for Denmark, Latvia and Romania, with many approaching 75% (Figure 4 and Figure S4b). Particularly high counts of several thousand small patches are found in Belgium, Germany, Finland, France, Hungary, Italy and Sweden. In Greece, Portugal and to a lesser extent Romania and Latvia, the size distribution is relatively flat, compared to other countries, with larger patches being similarly frequent as smaller ones.

**Figure 4.**
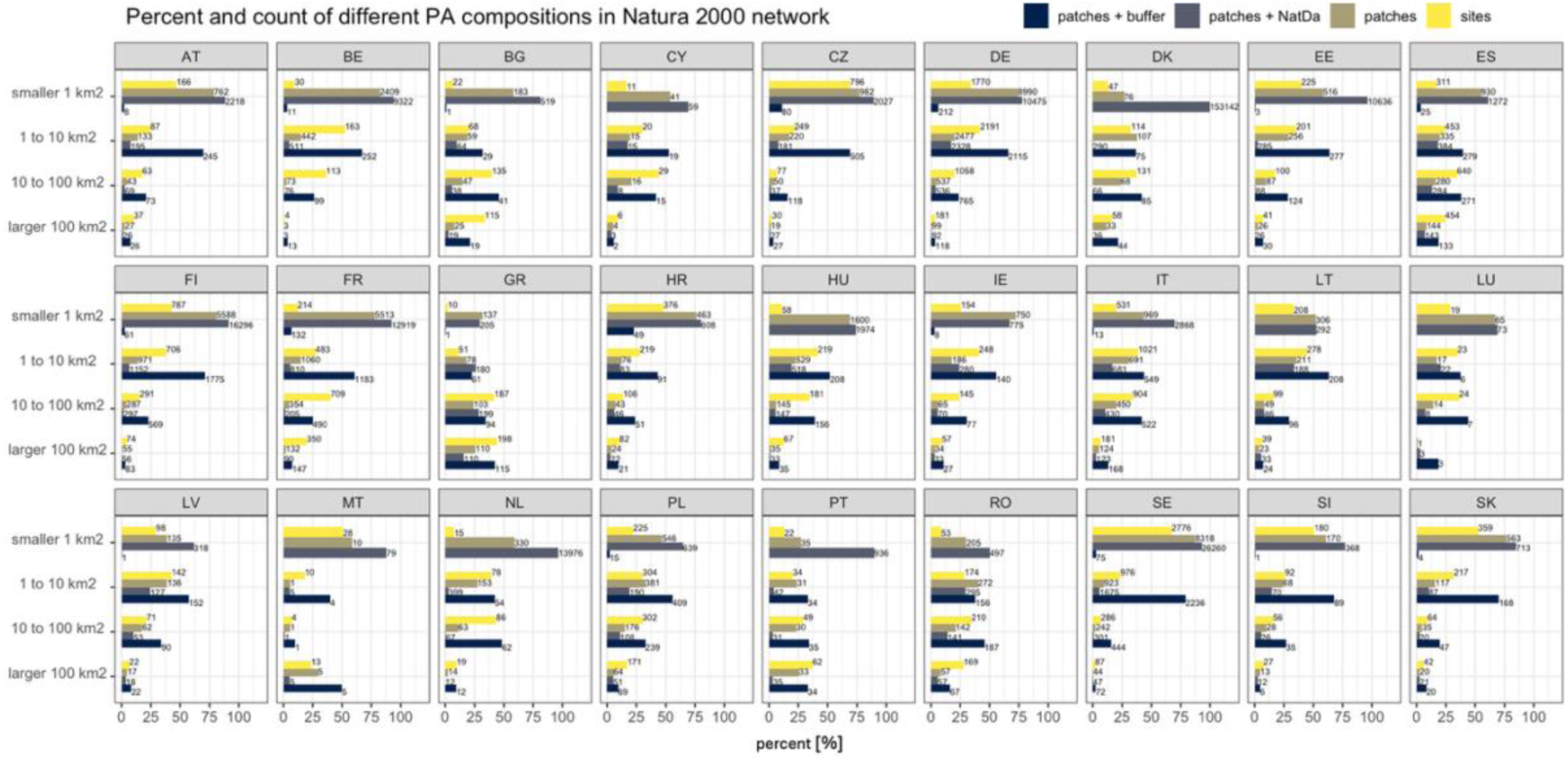
percentage (x axis) and count of different size classes (labels on bars) of protected area compositions sites, patches, patches plus national designations and patches plus 500-meter buffer in all Member States. Considering patches instead of sites shows a strong bias towards the smallest size group in almost all countries. Considering national designations in combination with Natura 2000 patches adds a substantial number in the smallest size category in most countries and strongly increases the bias towards very small patches in Denmark, Estonia, the Netherlands and Portugal. A buffer of 500m reduces the number of the smallest size category almost completely in most countries, with only few patches smaller than 1km^2^remaining. The reduction of numbers includes the larger size categories, showing that the close distance to the next patch is not only true for the smallest patches.

Considering nationally designated areas in the analysis decreases the amount of patches smaller than 1km^2^ only in Lithuania. The decrease in Lithuania is likely caused by nationally designated areas covering smaller Natura 2000 sites within their borders and that generally national designations are of larger size in this country. In Cyprus, Greece, Ireland, Latvia, Luxemburg, Malta and Poland the number of additional patches smaller than 1 km^2^ is less than 100, in all other countries significantly higher, with more than 150.000 additional patches in Denmark, and over 10.000 additional patches in Finland, the Netherlands and Sweden (Figure 4 and Figure S4c). The number of additional patches does not correlate with an increase or decrease in percentage of the size categories. The only country in which the percentage of the smallest size category is not dominant at the patch level with or without national designations is Greece. The highest increase of percent of small patches through consideration of national designations are found in Denmark (from 27% to almost 100%) and Portugal (from 27% to 90%), followed by Estland (58% to 96%) and Netherlands (59% to 97%). In Austria, Germany, Spain, Greece, Croatia, Hungary, Ireland, Lithuania, Luxemburg and Sweden the change in percentage of smallest patches is less than 10%. In most countries, the percent of national designations within the 500-meter buffer is less than 10%, and only Cyprus, Germany and Poland having more than a third of the 500-meter buffer area under national designations (Supplementary Material Table S3).

A buffer area of 500 meter around each patch eliminates the relative dominance of the smallest size group in all countries, as the distances from each smallest patch to the next patch are mainly short (Figure 4, Figure S1 and S4d).

Some countries have more isolated small patches in the landscape than others. Croatia retains 23% isolated smallest patches after buffering, followed by 10% in the Czech Republic, and ∼6% in France and Germany, while all other countries see a reduction to under 4% or complete elimination of the smallest size group.

### Summed area under each size class

While the vast majority of individual designations are of very small size, the vast majority of the European Protected Area extent is accounted by relatively few large protected areas, as it can be seen by plotting the cumulative protected area extent, starting from the smallest site first. The same pattern can be observed when repeating the analyses with individual protected patches and buffered patches (Figure 5 Error**! Reference source not found.**). The pattern is consistently observed across all Member States.

**Figure 5.**
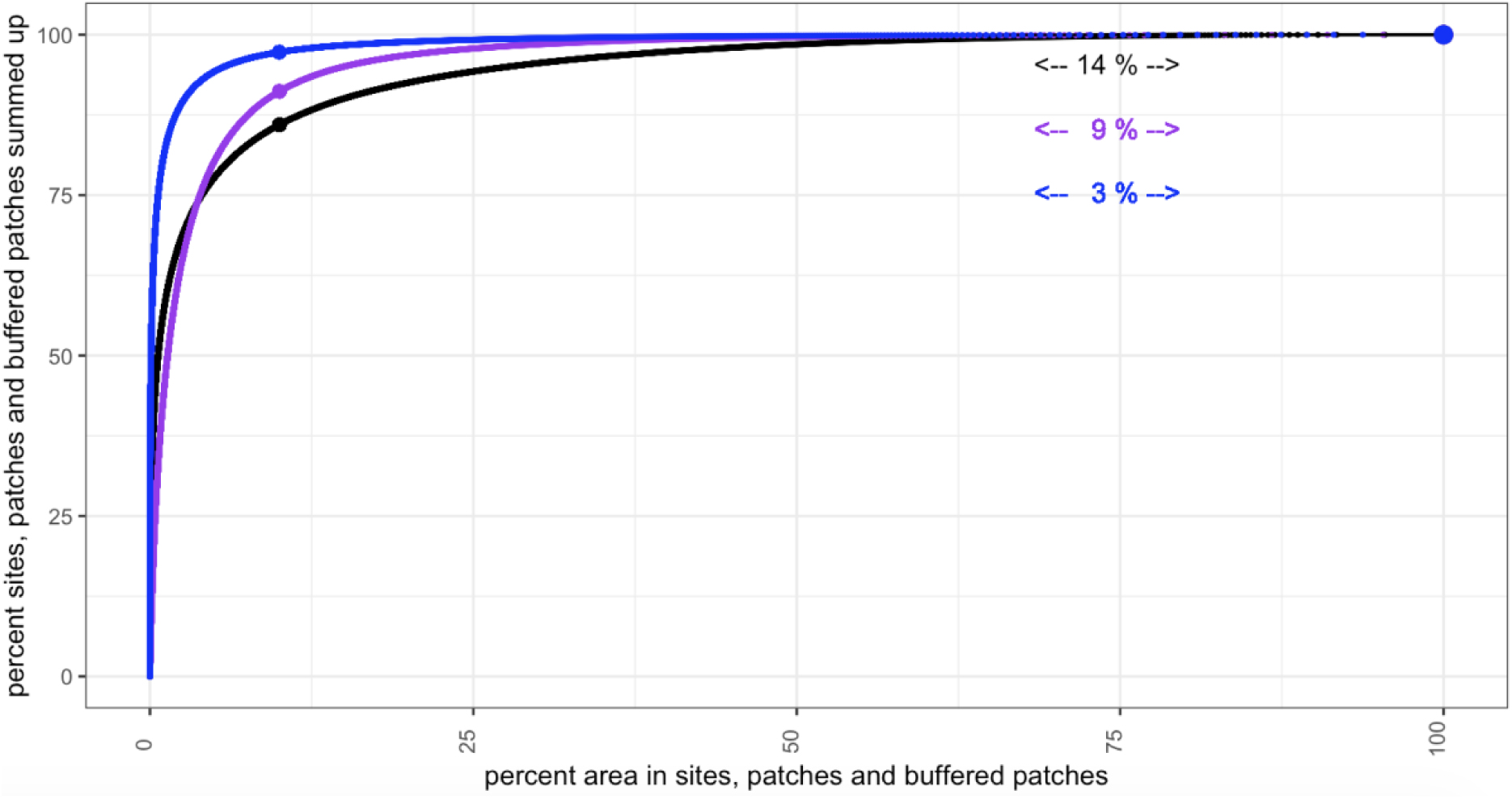
The cumulative sum of area of sites (black), patches (blue) and buffered patches (purple) in the EU, starting with the smallest sizes at the left. All curves share the general pattern that there is a sharp initial increase, indicating the large amount of individual spatial units with small size. The first point along the x-axis demarcates when 10% of the area is summed up. At the site level, this point is reached when 86% of all sites are summed up, the remaining 14% of sites (labelled under the curve) add up to 90% of the total area. At the patch level, 97% of patches add up to 10% of the area, and only 3% of patches contribute the remaining 90% of area. In all panels, the largest sites and patches are visible as distinct points at the end of the curve, each consecutive point contributing more area than the previous one. In each curve, the 3 largest sites or patches contain more than 10% of the total area.

At the EU level, 90% of the protected area extent is composed of the 4523 sites that are >30.5 km^2^, and these constitute only 14% of the total number of sites. This suggests that the vast majority of Natura 2000 sites contribute little to the total extent of the Natura 2000 network. The results are even more extreme when looking at patches, rather than sites. Specifically, 90% of the protected area extent consists of only 2405 patches that are >26.7 km^2^, and these constitute only 3% of the total number of patches. The smallest patch size starts at 5 m^2^, the median patch size is 0.165 km^2^, mean patch size is 22 km^2^ and maximum patch size 76.131 km^2^. This disproportion would be somewhat mitigated if each patch were expanded by a radius of 500 meters, in fact, the results are then similar to the site level, with 2567 patches (9% of the total number) representing 90% of the area extent, all being larger than 38.1 km^2^, however, with a much larger relative contribution of many of the smaller patches (Figure 5). Adding up all area documented for all sites in the Natura 2000 dataset leads to double counting of overlapping area at the site level and overestimates the total area by 200.000 km^2^ compared to analysis at the patch level.

### Compactness of Natura 2000 patches

The roundness metric, which measures the ratio between area and outline of any two-dimensional body. A perfect circle has a roundness of 1, while a completely flat polygon has a roundness of 0. All size categories have multiple patches with extreme values, indicating the existence of extremely high as well as extremely low compactness (Figure 6a). In all size categories, patches with almost perfect compactness exist at the maximum value of compactness, with decreasing roundness for values along the first quartile, median, third quartile, mean and maximum (Figure 6b). Patches at the third quartile of roundness appear to have a distinct core area across all size classes, and this is also the case for patches of median circularity in the largest size class. However, for all other size classes and lower roundness (mean, first quartile and minimum value of the metric), the shape and size suggest that we cannot distinguish a core area, or a core area exist only for a part of the patch. In other words, the distance from most points within a protected patch and the edge tends to be small, either because of the size of the patch, or because of its irregular shape or because of both. Example patches for minimal values show likely rivers and creeks within designated areas that do not have a buffer area along the riparian zone (Figure 6b, column min).

**Figure 6.**
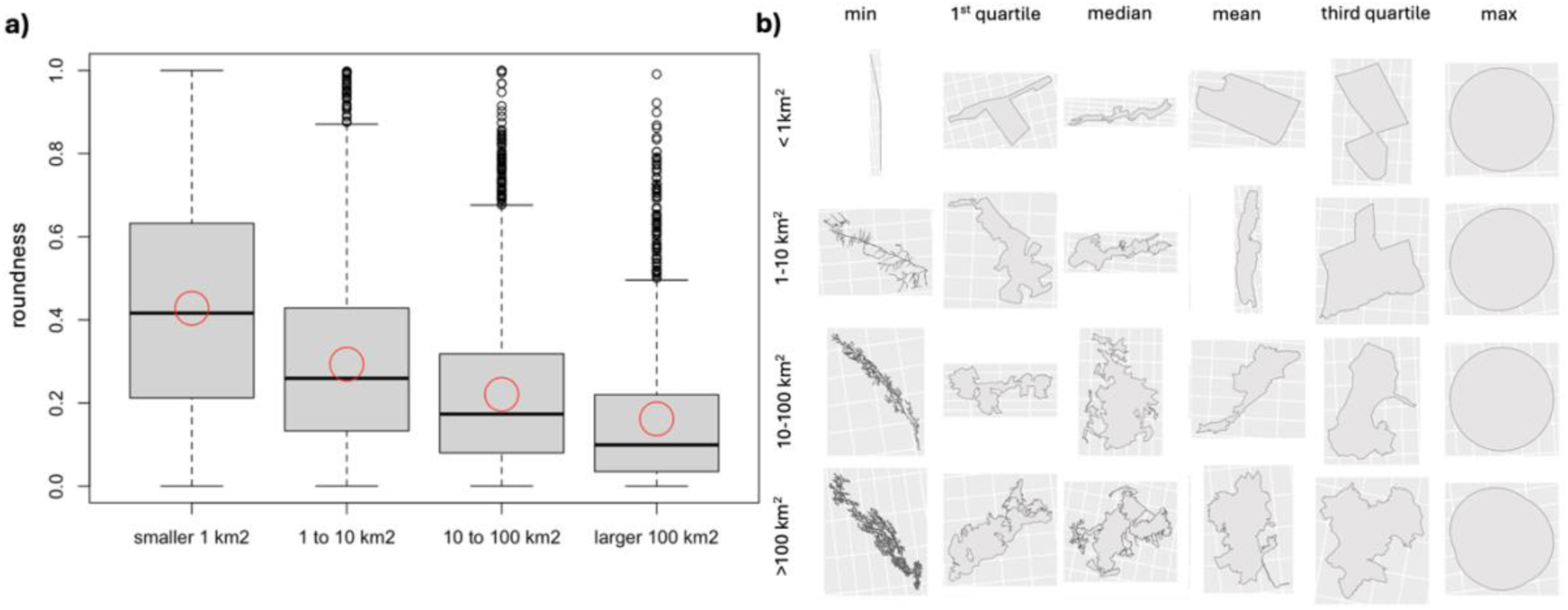
The measure of roundness in the four size categories of patches of Natura 2000 sites. Mean values depicted by the red circle. All categories include extremely high and extremely low values, but high values are increasingly rare in the higher size categories (Figure 6a). Smaller patches tend to be more compact, and three quarters of large patches are within the same range as the quartile with the lowest values in the smallest group. In the three larger size groups, values closer to 1 are exclusively outliers. The measure of roundness can be used as an indicator for compactness in all four groups, with almost perfectly compact patches at the maximum, relatively compact areas around the third quartile and mean, and increasingly complex and elongated forms above the mean, with almost linear forms at the minimum (Figure 6b).

### Distribution of size classes’ fragmentation across the European Union

Mapping the different size categories of Natura 2000 patches across Europe (Figure 7) shows strong clustering of patches smaller than 1 km^2^ in the Atlantic region in Belgium, high density in the Continental region in Germany and Eastern Europe, and high densities across Sweden and Finland. The largest patches seem to be evenly distributed across countries, with some gaps in the Atlantic region and the South of Finland and Sweden.

**Figure 7.**
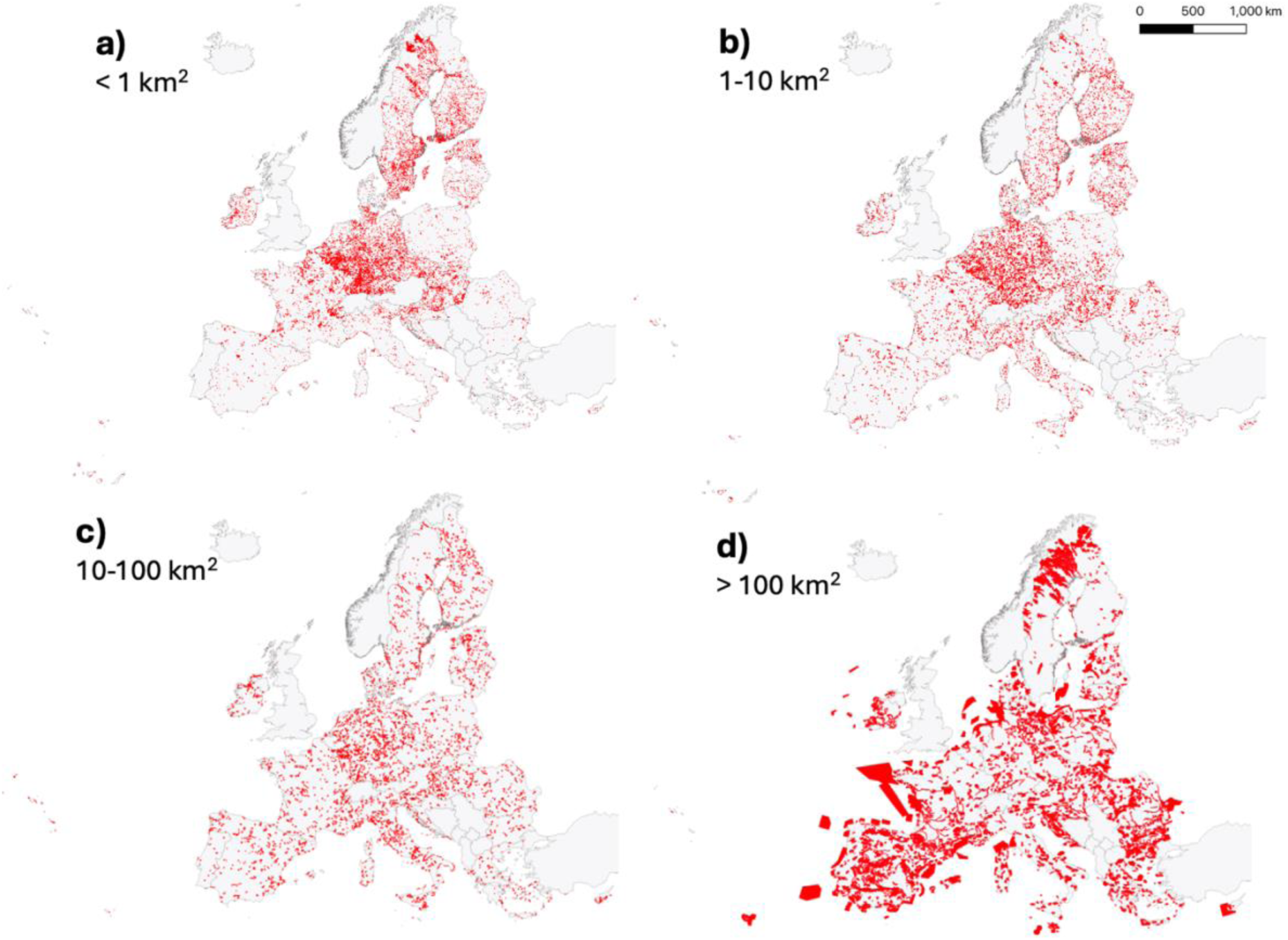
The location of each size category of Natura 2000 protected areas across Europe shows a clustering of the smallest size category of patches in South and North of Sweden, South of Finland, Estonia, South of Germany, Belgium, the Czech Republic, Slovenia, and Slovakia. Protected area patches between 1 and 10 km^2^ show a similar but less pronounced spatial pattern. Protected area patches between 10 and 100 km seem to be the most evenly distributed size category in Europe. The largest patches show some gaps in coverage in Northern France, Belgium, the Netherlands and Southern Sweden and Finland, with marine protected areas and Northern Finland being the largest.

### Land cover types in the EU

#### Land cover classification within and around Natura 2000 sites at EU level

Across Europe, forests account for the largest share within Natura 2000 sites (47%), followed by grasslands (27%). Other natural and semi-natural land cover types add up to 18%, 8% contain agricultural areas, and less than 1% are sealed surfaces. (Figure 8a). In all size categories of Natura 2000 patches and the 500 meter around them, forests cover the largest area, followed by grasslands. Compared to the two smaller size categories of Natura 2000 patches, the two larger size categories contain ∼4 % less forest and ∼3 % more natural systems other than grassland or forest. Within protected areas, other natural and semi-natural land cover types are the third largest category, while agricultural areas take up more area in the 500 meter around protected areas.

**Figure 8.**
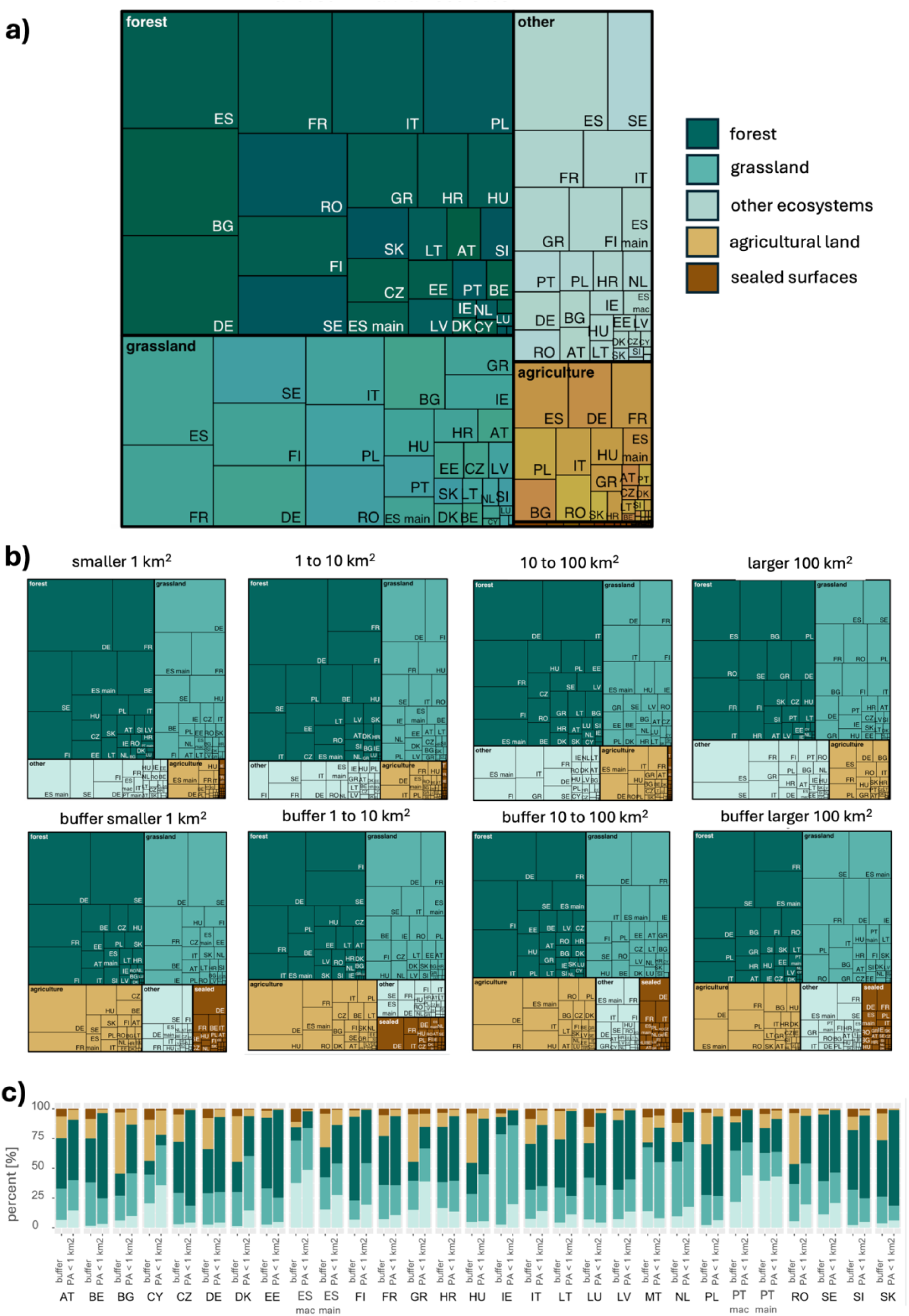
Relative cover of five land use and landcover categories forest, grassland, other ecosystems, sealed surfaces and agriculture, subset for each Member State’s share within Natura 2000 patches across the European Union (a). Patches and buffer areas are shown individually for each size category in (b). All size categories have forests as the largest share, followed by grasslands, both within and around protected area patches. Other ecosystems are the third largest group within Natura 2000 patches, and agricultural areas are the third largest group around Natura 2000 patches. Figure 8c shows paired bargraphs for each Member state showing the percentage of each landcover category in the smallest patches and the area within 500 meter around them. Results for Spain and Portugal are split between the mainland (main) and Macaronesian islands (mac).

While sealed surfaces are nearly absent within protected areas, they cover a similar area as ecosystems other than forest and grasslands the 500 meter around protected areas in all size categories (Figure 8b). In the Natura 2000 patches of the two larger size categories, the relative area of agricultural land is visibly larger than in the two smaller size categories, while this difference is not observed in the 500 meters surrounding the protected areas.

In the smallest size category, the increase of the combined area of sealed and agricultural areas around protected patches is ubiquitous across the EU, however, it varies in different magnitude across different countries, covering 25% and more of the 500 meter around Natura 2000 sites in Bulgaria, Cyprus, Germany, Denmark, mainland Spain, Greece, Hungary and Romania (Figure 8c). Portugal, Cyprus, Spain and Greece stand out as countries with the most natural and semi-natural land cover types other than forest and grassland in the Natura 2000 sites. Ireland, Malta and the Netherlands stand out with the most grassland.

### Species case studies

The protected area sites in which the three example species are reported in show varying degrees of internal fragmentation into separate patches (Figure 9). All species are found in countries where sites and patches are found in similar size categories, as well as countries where many more small patches exist than an analysis at the site level suggests. Visual inspection in a GIS viewer suggests that the analysis can severely underestimate the existing fragmentation of habitat available to species. For example, the little bustard, *Tetrax tetrax,* is recorded in Italy in the Natura 2000 site Costa Otranto - Santa Maria di Leuca. This particular site is listed with an area of 18.7 km^2^, and splitting sites into individual patches does not change the numbers in Italy much (Figure 9).

**Figure 9.**
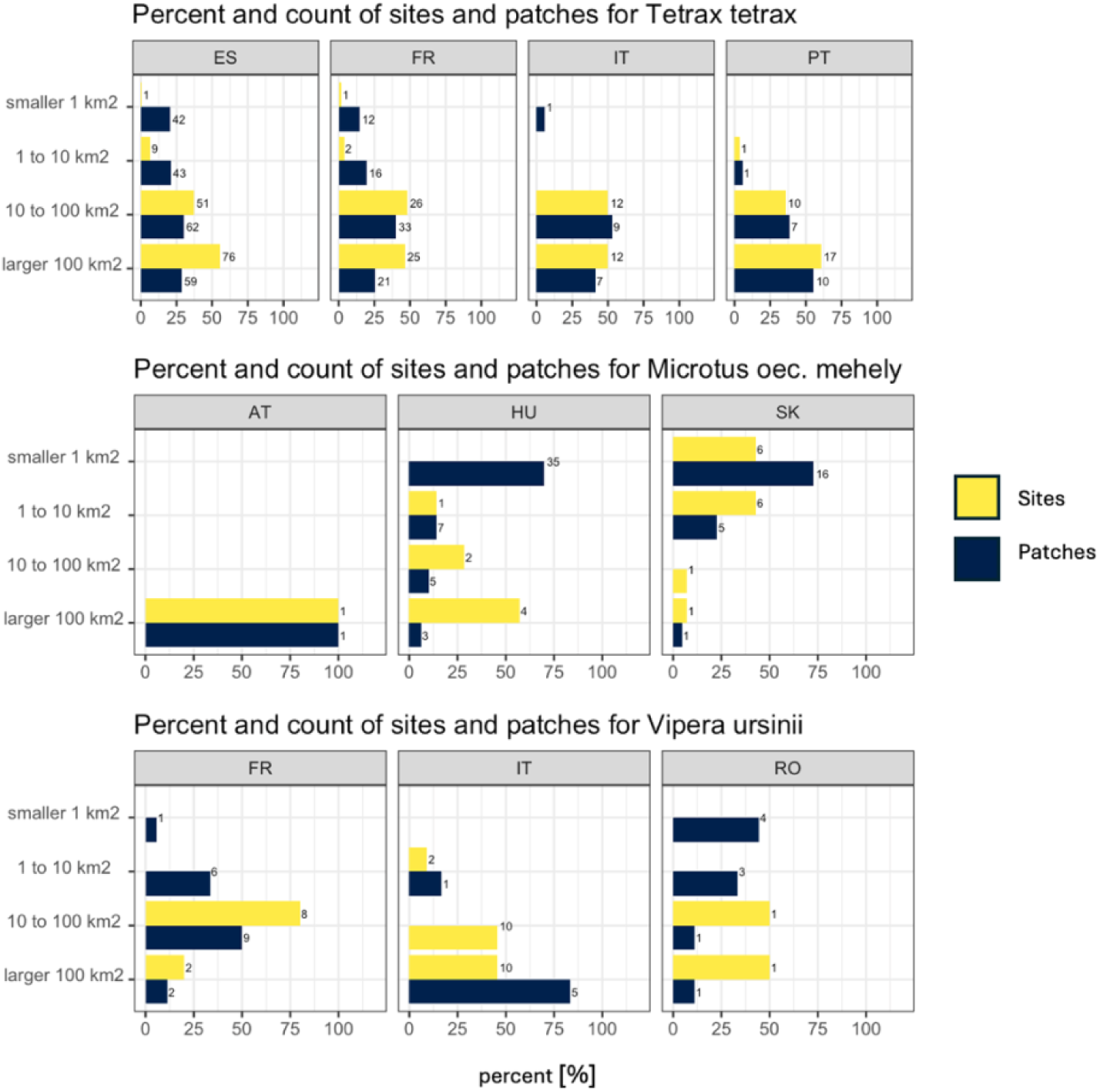
Count and percent of sites and patches in all countries of occurrence for the three examples species. All species are found in countries in which the sites they occur in include patches of much smaller size categories, as well as countries in which patches and sites show a similar relative composition of size categories.

However, visual inspection of the site shows that the little bustard has 28 smaller habitat patches available, surrounded by unprotected land, scattered along the coastline next to a major road. The analysis of sites and patches does not show this, as the site includes a large part of the ocean, and the fragmentation only becomes visible when only the terrestrial part of the site is considered. In contrast to that, several other larger sites in Italy overlap or share borders, which results in a larger connected and protected area than analysis at the site level would suggest. For example, the second largest site, Promontorio del Gargano, is listed with 700 km^2^. Because of overlaps with another 140 km^2^ large site the real area of connected and protected area is 830 km^2^ at the patch level.

In the case of the Pannonian tundra vole, the single large site in Austria turns out not to be fragmented, but visual inspection shows that the site encompasses a large lake with substantial amounts of build-up and agricultural area, suggesting that available habitat within the site is likely fragmented. Several larger sites in Hungary include 35 very small patches (Figure 9). In contrast to that, the sites in which the meadow viper is recorded in Italy overlap to a large extent, which means that there is a shift to a larger size category for patches compared to sites. In France and Romania, the two larger size categories contain equally numbers of sites and patches, but both smaller size categories exist for patches. A co-benefit of the analysis is the detection of likely errors in the official Natura 2000 database due to spelling mistakes. Extracted data from the Natura 2000 viewer included low numbers of records for *Tetrax tetrax* for Bulgaria, Greece, Latvia and Slovakia, where the species is officially extinct. These entries might be intended for the genus Tetrao, which would need to be verified with authorities.

## Discussion

### Size distribution of protected terrestrial Natura 2000 patches

Our analysis shows for the first time the size distribution of protected terrestrial patches within the Natura 2000 network, finding that on average every Natura 2000 site is made of four disjoint protected area polygons, and that ∼74% of the patches are ≤ 1 km^2^ and 92% are ≤ 10 km^2^. This pattern is generally consistent across countries. Nationally designated areas do not necessarily overlap with Natura 2000 sites, and the latter are often entirely outside national designations; considering all designations together shows that nationally designated areas cover in most countries less than 10% of the area within 500 meters around Natura 2000 sites, and add over 200.000 isolated patches smaller than 1km^2^ to the Natura 2000 network at the EU level.

### Isolation of protected terrestrial Natura 2000 patches

We found that the average distance between two Natura 2000 protected patches is 1 km, and 70% of the patches (n patches) are < 1 km apart. In fact, if each Natura 2000 patch were buffered by 500 meters, these patches would coalesce, bringing the total number down to 18.154 patches. These findings are broadly consistent across countries.

### Land cover within and around Natura 2000 patches

At the EU level, the dominant land cover within Natura 2000 sites is forest, occupying 46% of the Natura 2000 terrestrial extent, followed by grassland (28%), other natural and semi-natural habitats (18%, including wetlands, rivers, lakes, low-growing woody vegetation, mosses and lichens, sparsely vegetated and bare areas), sealed surface (<1%) and agriculture (8%). The differences to the buffer areas at EU level are clear but small, with increased agricultural and sealed surfaces of an additional 5% each, and reduced cover of other natural and semi-natural land cover types. However, there are stark differences across countries, for example, smallest patches are in multiple Member States surrounded by artificial land cover types between 20 and 50%.

### Implications for conservation

The extinction risk caused by land-based pressures outside of protected areas is thought to be higher than causes related to general habitat availability or habitat loss, making the consideration of matrix effects on biodiversity and the increased vulnerability of particularly small patches crucial for an effective improvement of conservation outcomes (Fletcher et al., 2024; Laurance et al., 2012; Ramírez-Delgado et al., 2022; Ries et al., 2017; Zabala et al., 2024). In this context, our results highlight two urgent priority areas for conservation planning, reporting, implementation and monitoring. One is management of edge effects at the local level to improve population viability at the patch level, and one is management of the matrix at the landscape level to improve metapopulation dynamics and enable range shifts under future conditions. Both are currently not considered in assessments of protected area effectiveness, despite their ecological significance.

### Local population viability depends on size and quality of available habitat at the patch level

Our results underscore the need to increase patch size and geometric compactness to decrease exposure to stressors and threats at the edges and optimise the local viability of populations. It is important to note that small size does not make a site less valuable, but mainly more vulnerable, as small patches often contain irreplaceable biodiversity in comparison to larger patches, while there is not much distance between core area and the surrounding land cover matrix (Hoffmann et al., 2018; Lindenmayer, 2019; Wintle et al., 2019). In many areas of central Europe, small patches are all that remains of once larger ecosystems in otherwise heavily modified landscapes. In the European context, small size has been found to be a contributing factor to population declines within and outside Natura 2000 sites, particularly in remaining grassland habitat within urban areas (Grossmann et al., 2023; Kajzer-Bonk & Nowicki, 2022). The analysis at the species level shows that realised fragmentation for terrestrial biodiversity might be much higher in sites that span multiple realms. However, the current data quality requires individual visual inspection in a GIS software and local knowledge of the exact location where the species is found to understand the true level of fragmentation a species experiences.

Designation alone is less effective than targeted management for achieving conservation outcomes, so simply enlarging a patch might not be sufficient (Geldmann et al., 2019; Langhammer et al., 2024). Targeted interventions have been found to substantially improve conservation outcomes for pollinators, butterflies, and amphibians in small patches in Europe (Grossmann et al., 2023; Kajzer-Bonk & Nowicki, 2022; Landler et al., 2023). In line with these findings, recent studies have shown that, on average, Natura 2000 sites are not performing significantly better than ecologically equivalent unprotected sites in terms of waterbird population trends (Wauchope et al., 2022). Although Natura 2000 sites showed similar trends over time compared to equivalent unprotected sites, the study found a large variation in performance, with protected area management, and protected area size, being the primary factor in determining protected area performance. Specifically, larger protected areas and protected areas with waterbird-specific management plans perform substantially better than other protected areas or equivalent unprotected sites (Wauchope et al., 2022). This study on waterbird populations adds to the vast body of empirical evidence for the negative impacts of small size, isolation and fragmentation on animal species (Fletcher et al., 2024; Laurance et al., 2012; Ramírez-Delgado et al., 2022; Ries et al., 2017; Zabala et al., 2024). In summary, additional designations increase patch size, while restoration within Natura 2000 sites increases habitat extent and quality, and mitigates suboptimal geometry that exacerbates edge effects. Likely priority sites are small patches and those with the largest amount of degraded or artificial land-cover within and around the protected area patch.

### Landscape-level dynamics on metapopulation dynamics and migration

Restoration in the direct vicinity of protected areas can provide two different ecological functions. It can reduce the magnitude or frequency of disturbance inside protected patches by providing a buffer, for example filtering pollution, and can increase species’ mobility and survival outside of protected areas through improved structural and functional connectivity between protected areas (DeFries et al., 2010; Hansen & DeFries, 2007; Kubacka et al., 2022; Lee et al., 2025; Zabala et al., 2024). Both functions, buffer and connectivity, need bolstering due to existing pressures. Clear trends of increased rates of land cover change in the immediate vicinity of protected areas have been found in Europe, calling for better monitoring of buffer zones and tighter legal mechanisms to prevent deterioration in these areas (Hellwig et al., 2019). Therefore, targeted interventions like restoration or land use management adjustments in the matrix surrounding protected area patches are particularly relevant to improving conditions for species sensitive to low or deteriorating connectivity.

In addition to species traits data and their ecological needs, information on the land cover types within and around protected area patches, as presented in the results, can help to identify priority areas for intervention (Sahlean et al., 2020). In some countries, the smallest Natura 2000 sites are surrounded by over 20% agricultural areas and by 10% built-up areas, including road networks within a 500-meter radius. Increase of agricultural area within protected areas is a global issue, with stronger increases of agricultural areas inside of protected areas than outside for the Palearctic region, which includes the EU (Geldmann et al., 2019; Meng et al., 2023). Together, this information corroborates suggestions for a more holistic management at the landscape scale, such as strategic zoning of allowed land use types (DeFries et al., 2010). Research in the UK has found a link between the magnitude of species declines and higher cover of agricultural areas, but also a positive impact of agri-environmental schemes, indicating an effective pathway to utilise funding for targeted interventions in the matrix (Mancini et al., 2023; Redhead et al., 2022)

### Important additional aspects to consider when prioritising targeted inventions

Small size alone is not a sufficient indicator for low quality or effectiveness of protection. Ecological thresholds in natural habitat cover are highly dependent on the ecosystem, and even built-up urban area can provide significant amounts of habitat for threatened species (Ives et al., 2016). Therefore, a specific high or low cover of any land cover type does not automatically indicate a problem that needs solving. However, several studies point to a minimum amount of 30-40% of intact habitat in a landscape before abrupt ecological changes are observed (Shennan-Farpón et al., 2021). This means that in the absence of restoration actions, many of the smallest sites might not only be unable to sustain the species and ecological processes for which they are established, but they are also unlikely to function as stepping stones or as viable habitat for patchy populations due to landscape-scale pressures exerted by the inhospitable matrix by which they are surrounded. Targets for land cover types, as well as setting objectives and targets for restored area or metrics for quality, would need to be based on ecologically meaningful assumptions (Carwardine et al., 2009; Geldmann, 2026; Palomo et al., 2025).

Another important aspect to consider when prioritising measures targeting improved habitat size, quality, and connectivity are species traits, as requirements differ significantly across functional groups, including cases where fragmentation can be a safeguard against spread of disease, fire or invasive species (Belote et al., 2020; Gilarranz et al., 2017; Isaac et al., 2018; Keeley et al., 2025; Sahlean et al., 2020). Existing recommendations can further guide the prioritisation and navigation of trade-offs between increasing size or improving management within or around small protected area patches, with a suggested ranking of importance from quality of protected habitat, to size of protected habitat, to connectivity of protected areas (Isaac et al., 2018).

### Required financial support

Both management at local and landscape levels depend on adequate support through financial and normative instruments. Several emerging and well-tested avenues exist that could be exploited more effectively.

The first example is upcoming implementation of the Nature Restoration Regulation. We urge national authorities, research organisations, non-government organisations, and other entities involved in the National Restoration Plans and in the implementation of restoration actions to conduct analyses like these to identify candidate sites and patches for restoration based on the likelihood of reduced ecological functioning because they are too small, oddly shaped, or too fragmented by human land uses.

Two more examples come from Denmark and Finland, who have already introduced financial instruments for the strategic purchase and compensation of private landowners that can be used to expand the protected area estate around small sites, and to restore land within and outside them. Denmark intends to invest 5.76 billion Euros of public funding to create new forest, restore wetlands and meadows and establish new national parks. Private landowners can voluntarily access funding through this scheme, which is intended to also reduce water, soil and atmospheric pollution from agricultural activities and increase natural carbon sinks (Danish Ministry for the Green Transition 2026).

In Finland, the METSO program targeting private forests has invested 400 million euros through permanent conservation covenants, fixed-term nature reserves, and purchase of private forest land by the state (Batpurev et al., 2025). Accounting for size, fragmentation and isolation could strategically direct METSO funding towards a more outcome-oriented payment for ecosystem service scheme (Batpurev et al., 2025). The HELMI program targets mires and bogs, wetlands and grasslands (Finnish Ministry of the Environment, 2026).

Our results show that grasslands cover 28% of the Natura 2000 sites’ terrestrial area, as well as of their immediate surroundings, and are the most common land-cover within and around the smallest Natura 2000 sites in Denmark, Ireland, Malta, and the Netherlands. Grasslands are at the same time one of the most threatened ecosystems, and have a known high rate of private ownership. A targeted approach to involve interested landowners for agri-environmental schemes in this area might yield large returns of investment (Haensel et al., 2023; Hagemann et al., 2025; Scholtz & Twidwell, 2022).

### Caveats and future priorities

#### Spatial data technical issues

Borders of the official datasets of administrative boundaries, and Natura 2000 and other protected area designations often do not align, leading to issues with small overlaps and cutoffs of spatial polygons when layers are intersected or combined, and high uncertainty of ecological fragmentation at sites covering multiple realms. Without improving data quality regarding alignment of boundaries and more precise location of biodiversity within sites, ecologically meaningful analyses of conservation effectiveness and impact of fragmentation will be hard to realise.

#### Other caveats

We chose to ignore all individual patches below 5 m^2^ and use a buffer size of 500 meters based on some ecological considerations and visual checks of the data, but of course, other values could be chosen and explored. As our aim was to produce a first understanding of large scale patterns, and other research points to 1000 meters as a distance in which most land conversions can be detected, we believe the chosen distance is a useful trade-off between capturing known thresholds linked to increasing anthropogenic land cover types while also representing a distance within other Natura 2000 patches are likely to be found, to inform further research on candidate sites for restoration or other management interventions to reduce pressures on small patches (Orsi & Le Clec’h, 2023). Satellite data is always based on technical categorisation of detected wavelengths in the imagery and, therefore, subject to potential misclassification and grey areas between classes. We conducted sensitivity testing of the land use categories used in this study by comparing the dataset with an officially validated dataset that was available for part of some Member States. Based on a test in Austria, we could confirm that natural and semi-natural species-rich grasslands cannot be distinguished from pastures or other anthropogenically used or improved grasslands with marginal value for nature with satellite imagery. This could be an important priority for future mapping approaches based on manual delineation and verification. Furthermore, agriculture and forestry can exist within forest and other vegetation classes without being detected by the satellite. Finally, some technical decisions could affect the results, as there are always trade-offs in consistency across different rules. One example is the analysis at the country level, which might have led to some duplication of extracted land use categories in buffer zones overlapping borders, but allowed to produce results for consistent buffer sizes for all Member States.

#### Future priorities

Besides addressing spatial geoprocessing issues there are additional research questions that we believe is important to further explore. For instance, only 2% of all patches represent 90% of the total area of the Natura 2000 Network. It would be helpful to understand habitat types and species are well represented in these large patches, and which are missing or underrepresented. Another important aspect would be to assess small patches with low compactness for their suitability for restoration and potential for designated buffer zones or expansion.

In summary, our study, describing the geometry of EU protected area estate and land use cover within and around Natura 2000 patches, combined with previous empirical findings of the impacts of patch sizes and edge effects on protected populations, suggests that there is a great potential to improve the status and trends of species within Natura 2000 sites. Our results show that increasing the size of small, protected areas of 50-500m^2^ can bring down the level of fragmentation by a factor of 4, likely offering feasible and impactful management interventions at the local and regional level by formally protecting intervening (non)habitat, through re-designing boundaries, land acquisitions, and funding programs. Strategically targeting this expansion only where it is ecologically necessary and feasible would result in an even smaller area increase required, thus reducing opportunity costs and trade-offs with other land use demands (Chapman et al., 2025).

## Acknowledgments

The presented analysis in this manuscript is the direct outcome of a conversation with Frank Vassen, European commission, at the Biogeographical Seminar series in the Macaronesian region in 2024. We thank Frank Vassen for his interest and suggestion to explore the question of internal fragmentation of Natura 2000 sites in Europe. Without him, this paper would not exist. We thank Frank Vassen and Bruno Combal (European Commission) for their extended support, interest, and contributing their expert knowledge to multiple discussions about the workflow and results, as well as their invaluable contributions to framing and editing of the manuscript. The work was funded under the European Union’s Horizon Project “Natura Connect” under grant agreement number 101060429.

