## Supplementary Material S1-S6 for "Small patches dominate the Natura 2000 network of protected areas in Europe"

### Supplementary Material S1: Table S1: Data sources

Table S1: Data sources and processing steps

| **Data [original name]** | **Source URL and further description** |
| --- | --- |
| Natura 2000 sites  [Natura 2000_end2021_rev1_epsg3035.shp] | [https://www.eea.europa.eu/data-and-maps/data/natura-14](https://protect.checkpoint.com/v2/___https://www.eea.europa.eu/data-and-maps/data/natura-14___.YzJlOmlpYXNhOmM6bzo5NGNlZWYxMmMyZWM2NjI3YTU3ZmY0N2RiMmI2NGU2Mzo2OmIyYTM6ZTk4NWVkZGYyNTdiNTIzMTdkMTM1YTM5N2NlNWIwZGE2ZGZlZDJlMTQ5Zjc2MWNkMDdjZTdhYWJhMDE5YjNkNjpwOkY6Tg) |
| NatDa sites | <https://www.eea.europa.eu/en/datahub/datahubitem-view/f60cec02-6494-4d08-b12d-17a37012cb28?activeAccordion=1097586%2C1086849>  Sites were used as standalone dataset to calculate percentage of coverage in the buffer zones, and in combination with Natura 2000 sites in a second scenario to test the dependence of the level of fragmentation in the Natura 2000 network on national designations. |
| N2K_2018_3035_v010_fgdb | <https://land.copernicus.eu/en/products/n2k/n2k-2018#download>  Name : N2K Land Cover/Land Use 2018 (vector), Europe, 6-yearly  Manual: <https://library.land.copernicus.eu/products/N2K_2006-2018_PUM_v1.html>  Natura 2000 sites were selected to represent sites rich in grasslands. As grassland is likely one of the more difficult classifications due to limitation of management classification that a satellite image can produce, the dataset seemed suitable to check mismatches in accuracy between this and the less detailed CLC + Backbone data.  The data is a Copernicus Core service product. Core service name: Copernicus Land Monitoring Service, run by EEA. We used the most recent version, dated 2018.  To assess how accurate the land cover classes of the “CLC + backbone” raster data are, tabular statistics were done in areas of overlap between CLC and the fine resolution data for Austria to serve as sensitivity test. Histograms and flow diagrams were made for different numbers of categories and subcategories. |
| CLC + backbone (10m) | [https://doi.org/10.2909/b0bd43c6-1fa1-4d88-9c45-98b13a95d0b2](https://protect.checkpoint.com/v2/r02/___https://doi.org/10.2909/b0bd43c6-1fa1-4d88-9c45-98b13a95d0b2___.YzJlOmlpYXNhOmM6bzo0YzI5MTBjNDlkYWM0YjdkNzBlMWQwODE4M2QyOGE0YTo3OjM4ZDU6NDIxZjM1YjU2ZjU4ZTFmNWYyZTM3NjE2NGRjNjEzNmJjMWFlNjM1NGJjM2VkYzliMjFmNzJlYmI4NjIyYzBmOTpwOlQ6Tg) |
| admin boundaries | https://ec.europa.eu/eurostat/web/gisco/geodata/statistical-units/territorial-units-statistics |
| Buffer:  500m | For sites, it is important to keep site integrity for a site-level analysis. As many sites overlap with each other, assigning buffers to individual sites is not leading to any meaningful spatial polygon type. |
| Species’ occurrence at Natura 2000 sites | Searchterms of species name in the Natura2000 viewer and download of site IDs as csv file  https://natura2000.eea.europa.eu |

### Supplementary Material S2: Figure S2: distance to next patch

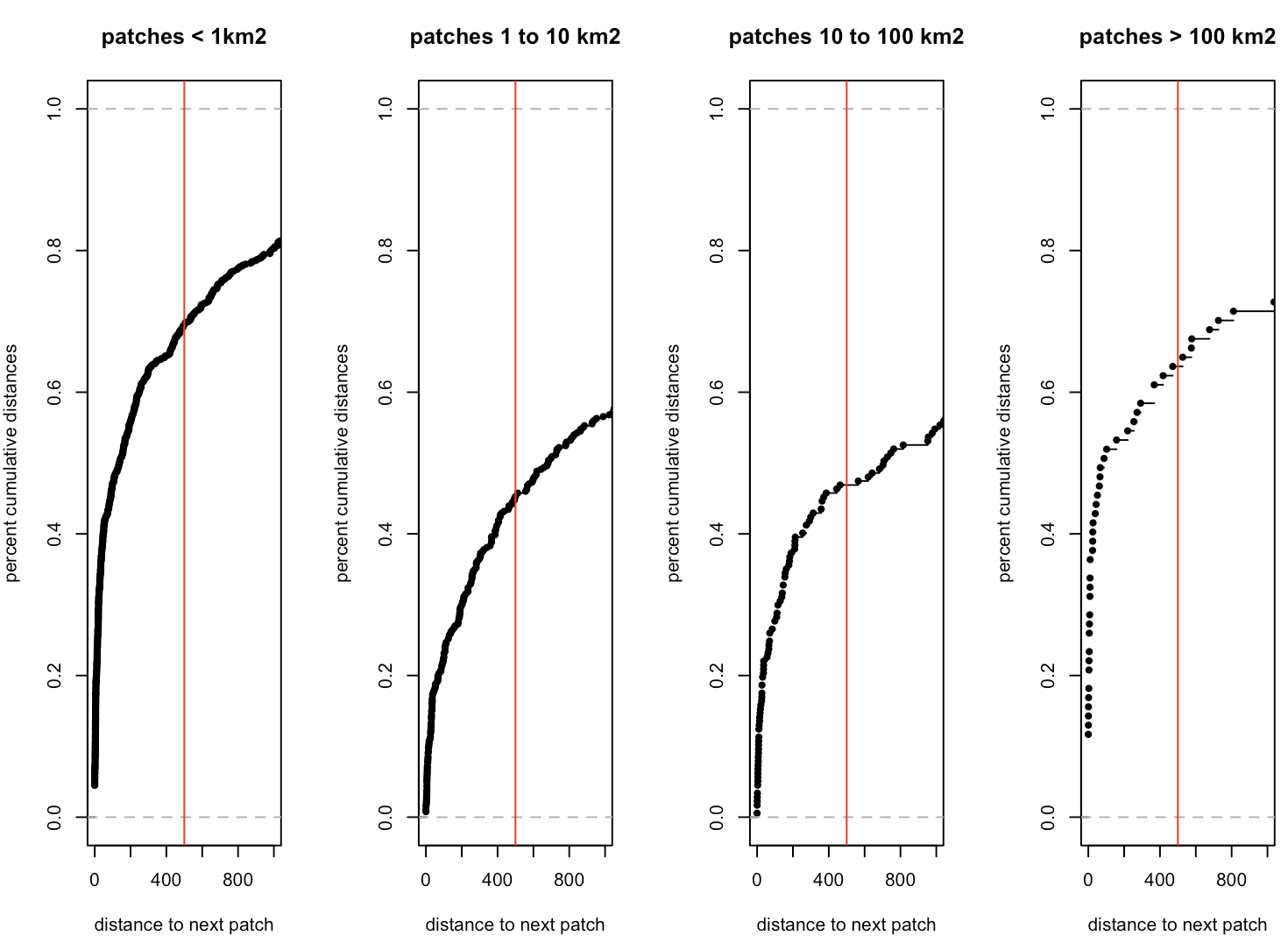

Figure S1: Cumulative distance to the next patch in meters for each size category. The red abline shows a distance of 500m, resulting in 70% of patches being connected to the next closest patch.

### Supplementary Material S3: fragmentation analysis of clipped Natura 2000 to terrestrial Europe

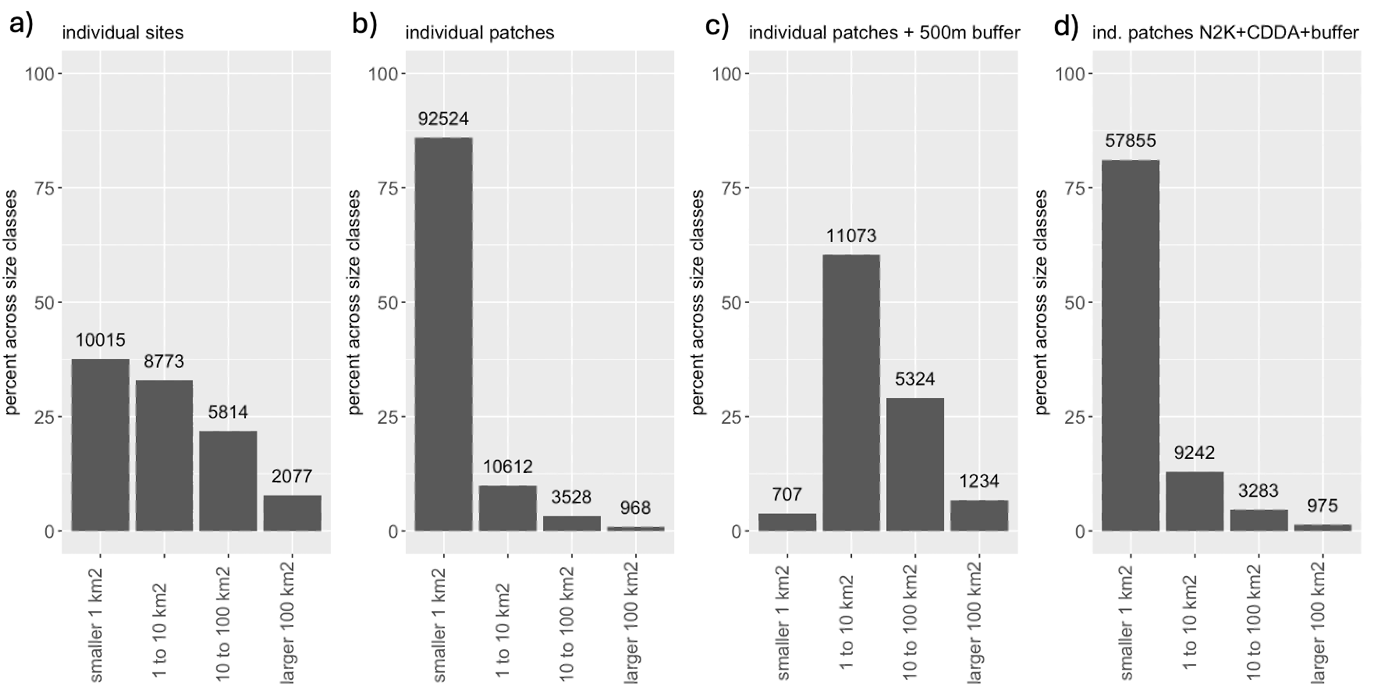

Figure S2: size distribution and count of sites and patches in different size categories using the dataset of clipped terrestrial Natura 2000 dataset.

### Supplementary Material S4: Percentage and count of size classes of different protected area compositions in European Member States

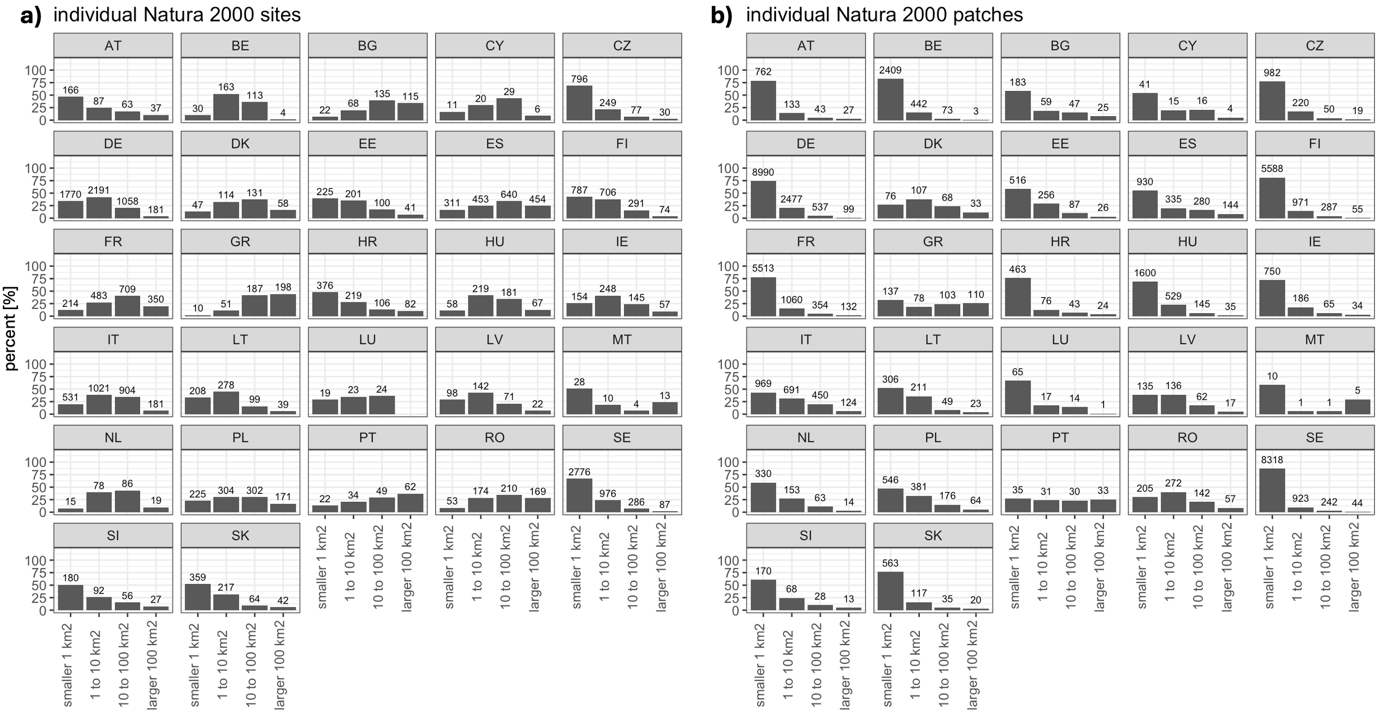

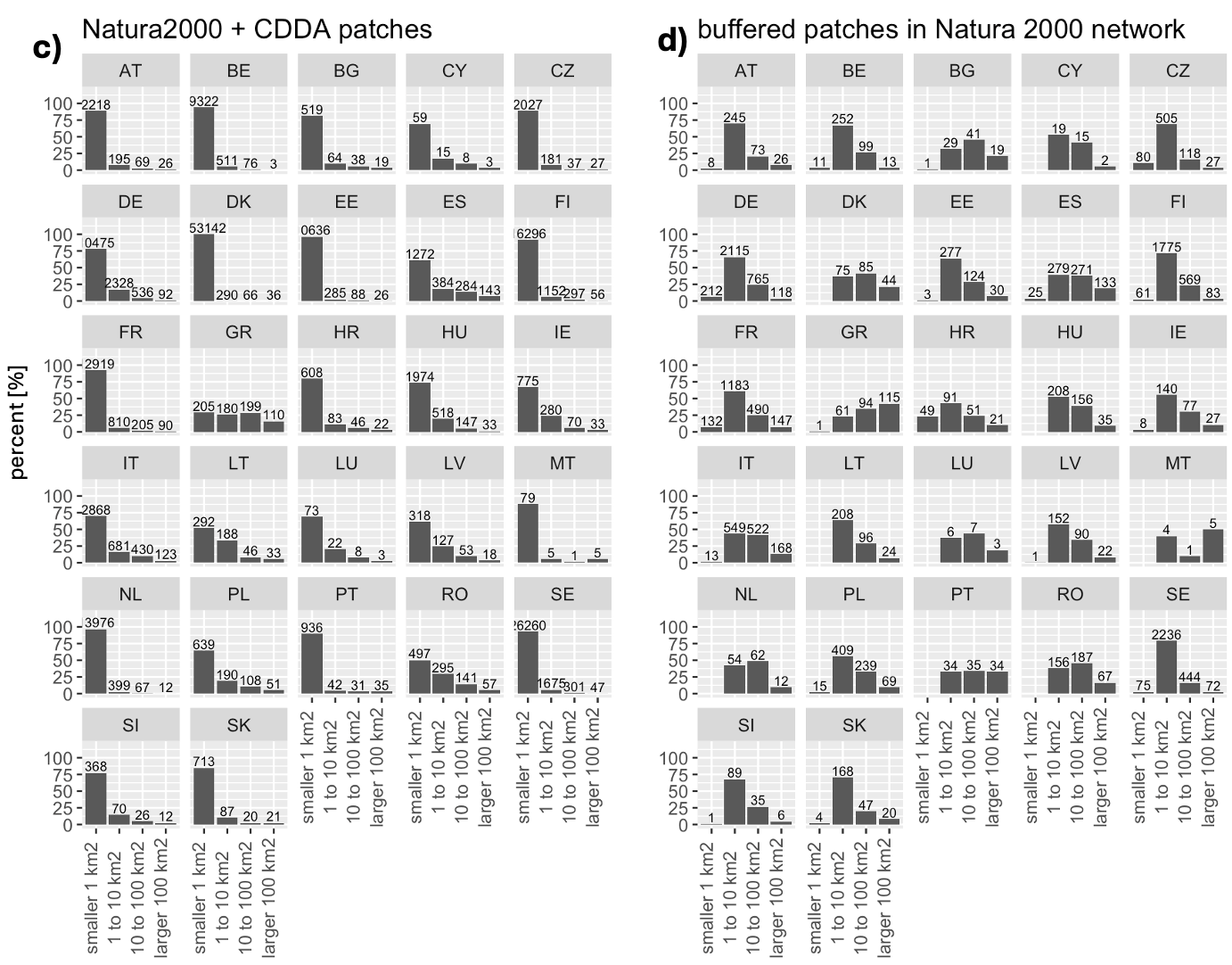

Figure S.1: percentage (y-axis) and count of different size classes (labels on bars) of sites(a) and patches (b) in each Member State. Considering other conservation area designations adds a substantial number of patches in most size categories, but reduces the bias towards very small areas in only a few countries (c). A buffer of 500m reduces the number of disconnected patches by about 80%, and the total number of patches is lower than the number of Natura 2000 sites, with only few patches smaller than 1km^2^. The reduction in numbers includes the larger size categories, showing that the proximity to the next patch is not limited to the smallest patches (d).

### Supplementary Material S5: fraction of National Designated Protected Areas within a 500-meter radius around Natura 2000 sites

Table S3: Calculated percentage of national designations within a 500-meter radius around Natura 2000 patches, sorted for increasing percentage from top to bottom.

| **Member State country code** | **National designations in 500-meter radius around Natura 2000 [km^2^]** | **Area of 500-meter buffer around Natura 2000 [km^2^]** | **Percent cover of national designations in a 500 m buffer** |
| --- | --- | --- | --- |
| IE | 73.1 | 11161.8 | 0.7 |
| RO | 276.7 | 18097.9 | 1.5 |
| MT | 10.2 | 498.0 | 2.0 |
| HU | 289.6 | 12281.4 | 2.4 |
| ES | 1220.0 | 47804.2 | 2.6 |
| BE | 198.8 | 6634.1 | 3.0 |
| HR | 246.0 | 7210.4 | 3.4 |
| SE | 2158.1 | 59375.6 | 3.6 |
| EE | 223.0 | 5933.8 | 3.8 |
| FI | 898.4 | 21899.2 | 4.1 |
| SI | 212.1 | 3877.7 | 5.5 |
| PT | 331.7 | 5663.3 | 5.9 |
| IT | 1815.6 | 26968.9 | 6.7 |
| LV | 277.3 | 3556.6 | 7.8 |
| DK | 453.5 | 5431.6 | 8.3 |
| BG | 1097.9 | 12350.7 | 8.9 |
| GR | 1311.5 | 11262.7 | 11.6 |
| NL | 482.3 | 3865.2 | 12.5 |
| LU | 105.0 | 821.3 | 12.8 |
| LT | 1130.1 | 6475.8 | 17.5 |
| SK | 1067.8 | 5544.8 | 19.3 |
| CZ | 1451.7 | 6906.9 | 21.0 |
| AT | 1571.3 | 6895.4 | 22.8 |
| FR | 14181.9 | 55501.3 | 25.6 |
| DE | 23871.2 | 66074.7 | 36.1 |
| PL | 8649.4 | 21624.1 | 40.0 |
| CY | 483.5 | 1124.8 | 43.0 |

### Supplementary Material S6: Accuracy of land use categories based on comparison of CLC + Backbone raster data with Copernicus Core Service product in vector format

The validated fine-resolution polygon data from the Copernicus Core Service has 8 broad categories, with several subcategories at the second, third and fourth level of increasing detail. The fourth level includes only a limited number of land use classes, and was not considered in the testing. The CLC+ Backbone satellite raster data has 14 categories. We used Austria as a case study to compare how well the different categories in the two datasets match each other. Results show a high match for several categories, with some caveats (Table S6, Figures S6.1 to S6.6). Out of the 14 categories, three are not relevant for land use and land cover (Coastal Seawater Buffer, Outside Area, and No Data), two were not present in Austria (Lichen and mosses and evergreen broadleaved trees), and the remaining nine were present in Austria to be tested (Table S6).

Table S6: land use and land cover categories in the two different datasets and likely matches

| **Vector layer Copernicus Core Service** | **10m raster CLC + Backbone** | **Test of matching** | **Match quality** |
| --- | --- | --- | --- |
| Urban + other buildup, including parks/leisure  Main category # 1  2^nd^ categories: 4  3^rd^ categories: 0 | Sealed | Sealed surfaces show several mismatches between the two datasets. While the “urban and infrastructure” category in the vector data also contains several other categories from the raster data, such as trees, sparsely vegetated areas or permanent herbaceous areas, the category of sealed surfaces in the raster data contains several other categories from the vector dataset, such as gare shared between both datasets. Therefore, some sealed surfaces are likely better reflected in the CLC-Backbone data and some better in the polygon data. For example, the polygon data includes fallen land, military, sports areas and parks in the urban category, which likely get more accurately classified as vegetation in the satellite data. On the contrary, especially road infrastructure seems to be difficult to detect with satellite imagery when it is covered by crowns of trees. Percentage-wise, the category for sealed surfaces in the satellite imagery is the only category that shows a large match with the urban category in the vector data. | ok |
| Agriculture, incl agroforestry  Main category # 2  2^nd^ categories: 3  3^rd^ categories: 4 | Periodically herbaceous | The absolute majority of agricultural land in the polygon data falls into this category in the raster data, likely because agricultural fields show up as vegetated only part of the time, and unvegetated after harvest until the next growing season. This category falls to a smaller extent into the sealed category of the raster data (likely fields close to sealed surfaces and built-up areas), and to an even smaller extent into the permanent herbaceous category (likely permanent crops). | good |
| Forest  Main category # 3  2^nd^ categories: 6  3^rd^ categories: 0 | 3 categories:  Needle leave,  Broadleaved,  Evergreen broadleaved | Forest in the CLC+Backbone dataset seems to consist almost exclusively of forest-related categories from the high-resolution polygon dataset. | good |
| Grasslands (managed and unmanaged)  Main category # 4  2^nd^ categories: 2  3^rd^ categories: 2 | Permanent herbaceous | Although most classified grassland in the polygon data is captured in the permanent herbaceous class in the raster layer, there are 25-30% of pixels that match other categories, mainly forest, but also agriculture, shrub and sparse vegetation from the polygon dataset. | medium |
| Heath and shrub  Main category # 5  2^nd^ categories: 3  3^rd^ categories: 0 | Low woody vegetation | Almost all of the classified heath and shrub in the polygon data is captured by the low woody vegetation category in the raster data. To a smaller extent, forests from the polygon data also fall into this raster category. | ok |
| Sparsely vegetated areas  Main category # 6  2^nd^ categories: 3  3^rd^ categories: 3 | 3 categories:  Non & sparsely vegetated area,  Lichen & mosses,  Snow & ice | Both non and sparsely vegetated areas, as well as snow and ice, almost exclusively match the sparsely vegetated areas in the polygon dataset. Lichen and mosses are not classified for Austria. About 10 % of the non and sparsely vegetated areas in the raster data match the forest, shrub, and grassland classification in the polygon dataset. | good |
| Marsh & peat and intertidal (wetlands)  Main category # 7  2^nd^ categories: 2  3^rd^ categories: 3 | Permanent herbaceous | This class does not exist in the CLC-backbone raster for Austria and could not be assessed |  |
| Water  Main category # 8  2^nd^ categories: 4  3^rd^ categories: 2 | Water | Water in the raster dataset seems to match almost exclusively the water-related categories from the high-resolution polygon dataset. | good |
|  | Coastal Seawater Buffer | Not present in Austria, therefore not tested | N/A |
|  | Outside area | Not present in Austria, therefore not tested | N/A |
|  | No Data | Not present in Austria, therefore not tested | N/A |

#### Details of testing the matching of both land cover datasets across categories and subcategories

The sensitivity analysis provided additional insights into the usefulness and caveats of satellite-derived land use classification data for spatial analysis of conservation planning and management. We matched the first-level classification from the polygon data to the raster data by counting the cells of each raster category within each polygon class, and then plotted the percentage as stacked bar plots and a flow diagram (Figure S6.1 and S6.2). Of the 9 categories present in Austria, 6 showed a good match, with over 90% matching categories (Figure S6.1 and S6.2). For categories with more convoluted matching, where multiple categories in one dataset were captured by one category in the other dataset at a higher percentage, we also produced flow diagrams for subsets of the data using second- and third-level classification to identify potential reasons to explain the existing mismatch. Under the assumption that the main intention is to identify non-natural use types that might have negative impacts on biodiversity, it is a priority to have a clear identification of 1) built-up areas and 2) agricultural areas. Therefore, we examined matches for all non-natural categories in the polygon data to the raster data (Figure S6.3), matches from permanent and herbaceous land use in the raster to the polygon data (Figure S 6.4) and matches from sealed surfaces from the polygon data to the raster data (Figure S 6.5 and Figure S6.6).

The first investigation of matches between the second level of all anthropogenic subcategories in the polygon layer (urban and agriculture) shows a strong match between arable land

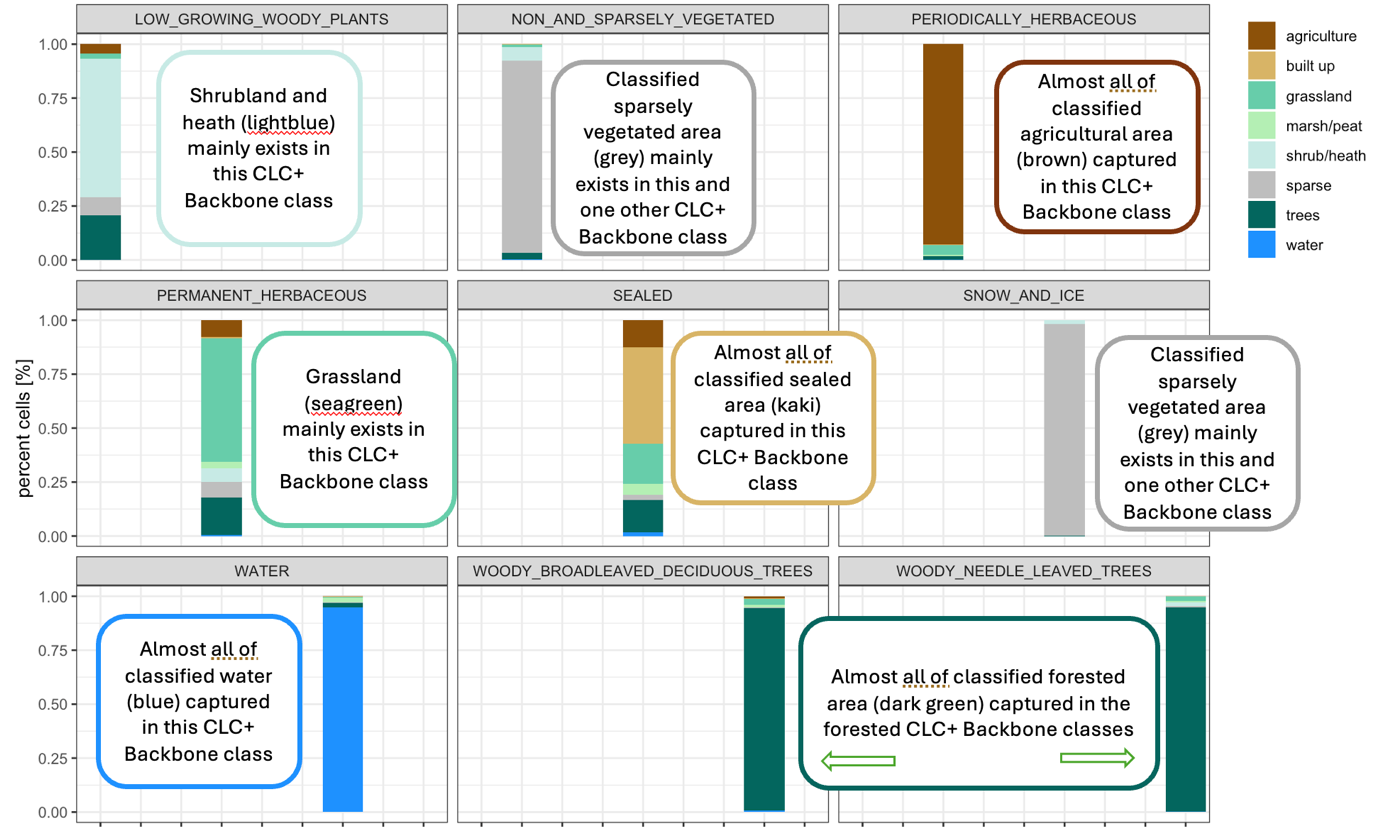

**Figure S 6.1:** Match between classification in the polygon data (colors) and the 10m raster data (panels). Panels show the share of different LULC classes from the polygon data within LULC types in the raster data. The color scheme shows LULC types in the polygon file. Data were tested for areas covered by the validated polygon data in Austria.

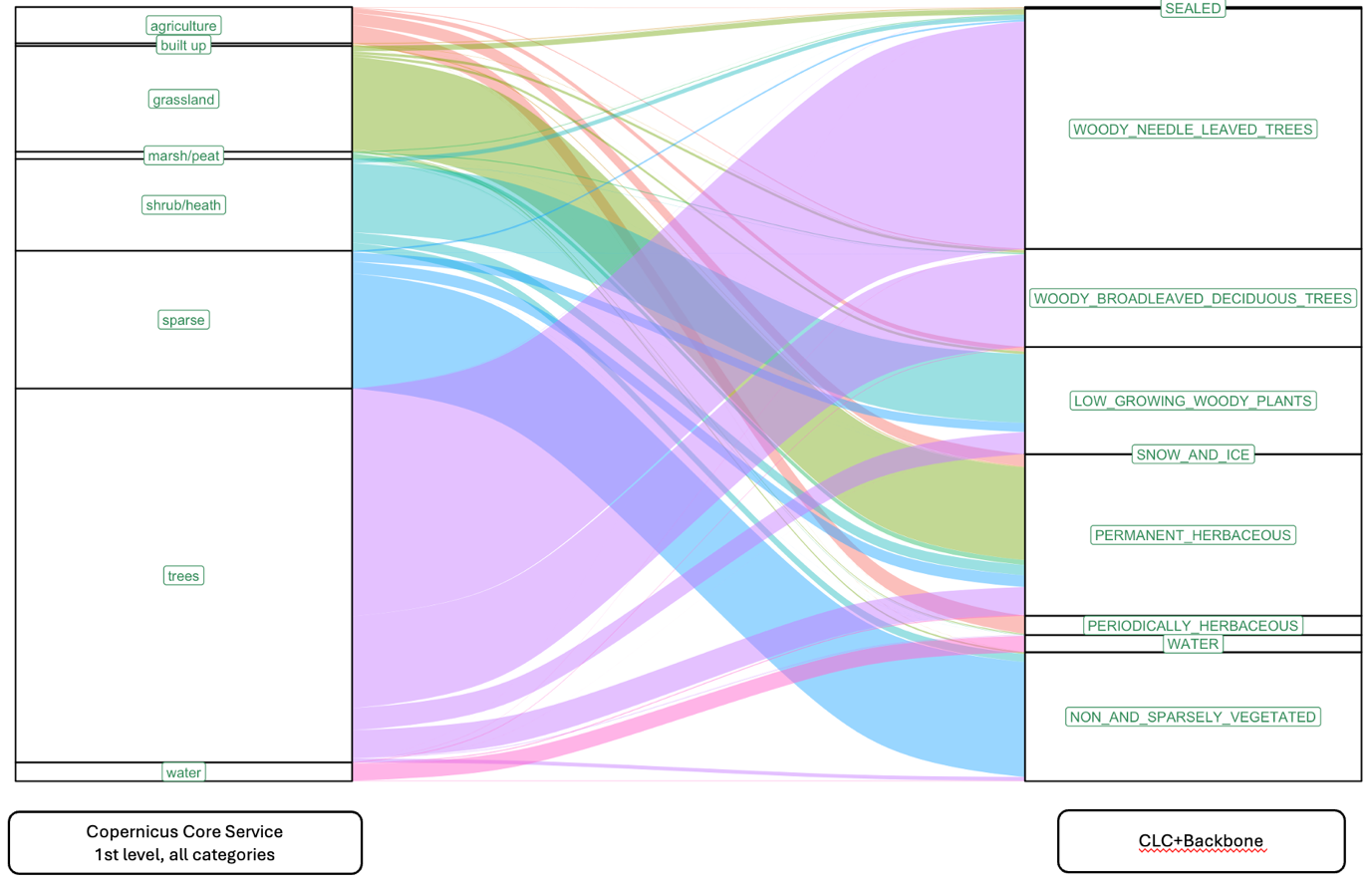

**Figure S6.2:** Flow between land cover classifications in the two datasets. Large matching flows exist for forests, sparse vegetation, and water. The match between “built-up” and “sealed” classifications is too small to assess in this visualisation. Agriculture flows from the polygon data into several natural land cover categories in the raster data, suggesting additional testing for these categories.

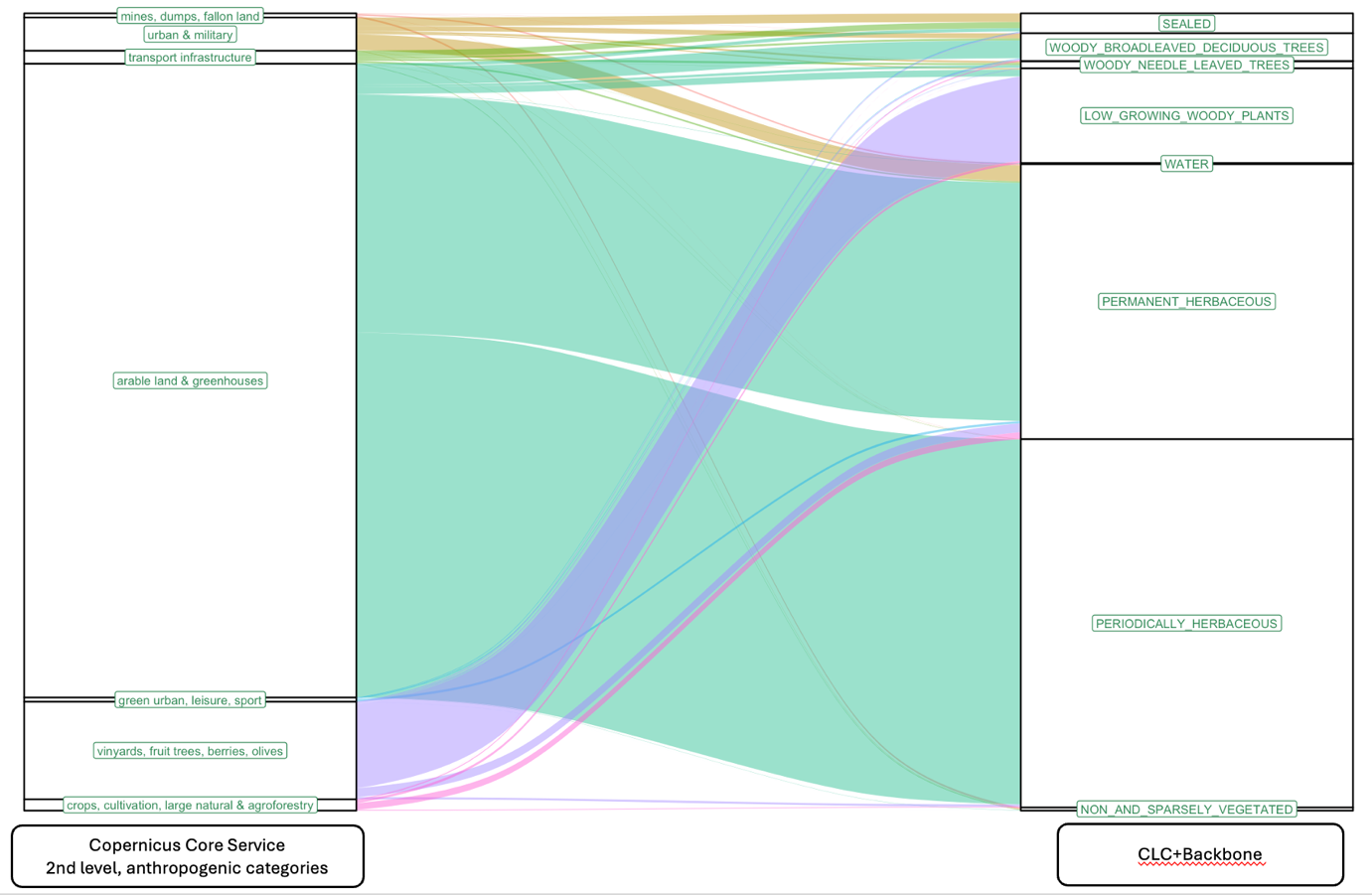

**Figure S6.3:** Making use of the 2nd level of classification with more detail in the polygon data for the subset of the two anthropogenic categories, urban and agriculture, shows that arable land is mainly captured in the raster categories “periodically herbaceous” and “permanently herbaceous”. Vineyards and berries are found to a large part in the raster category “small woody vegetation”, and traffic infrastructure is hidden to a large extent in the different tree categories.

(polygon) and periodically herbaceous (raster), indicating high certainty that the periodically herbaceous category captures almost exclusively agricultural land (Figure S6.3). However, another strong flow from arable land goes to permanent herbaceous, indicating that it is not possible to distinguish natural grasslands from potentially intensively used pasture lands. Another potentially high-intensity use type, such as berry farms and vineyards, shows a strong flow to the land cover category of low growing woody plants, indicating another potential mismatch between an anthropogenic and natural land cover type that makes it impossible to exclude the possibility of high use of pesticides and other disturbances in this category in the satellite imagery. A strong flow also exists from transport infrastructure to the forest classes in the raster layer, indicating that the satellite detects tree crowns above sealed road surfaces, concealing the pressure on biodiversity in this category (Figure S6.3). Reversing the flow from both herbaceous layers in the satellite data to the third level of subcategories in the polygon data shows a clearer split between the good match between the periodically herbaceous category and exclusively anthropogenic classes, and the mixed flows from the periodically herbaceous class to the natural as well as managed grasslands (Figure S6.4). Based on the testing, we characterise the permanent herbaceous category as the most difficult category, as it includes large flows between both natural and anthropogenic classifications in the polygon dataset.
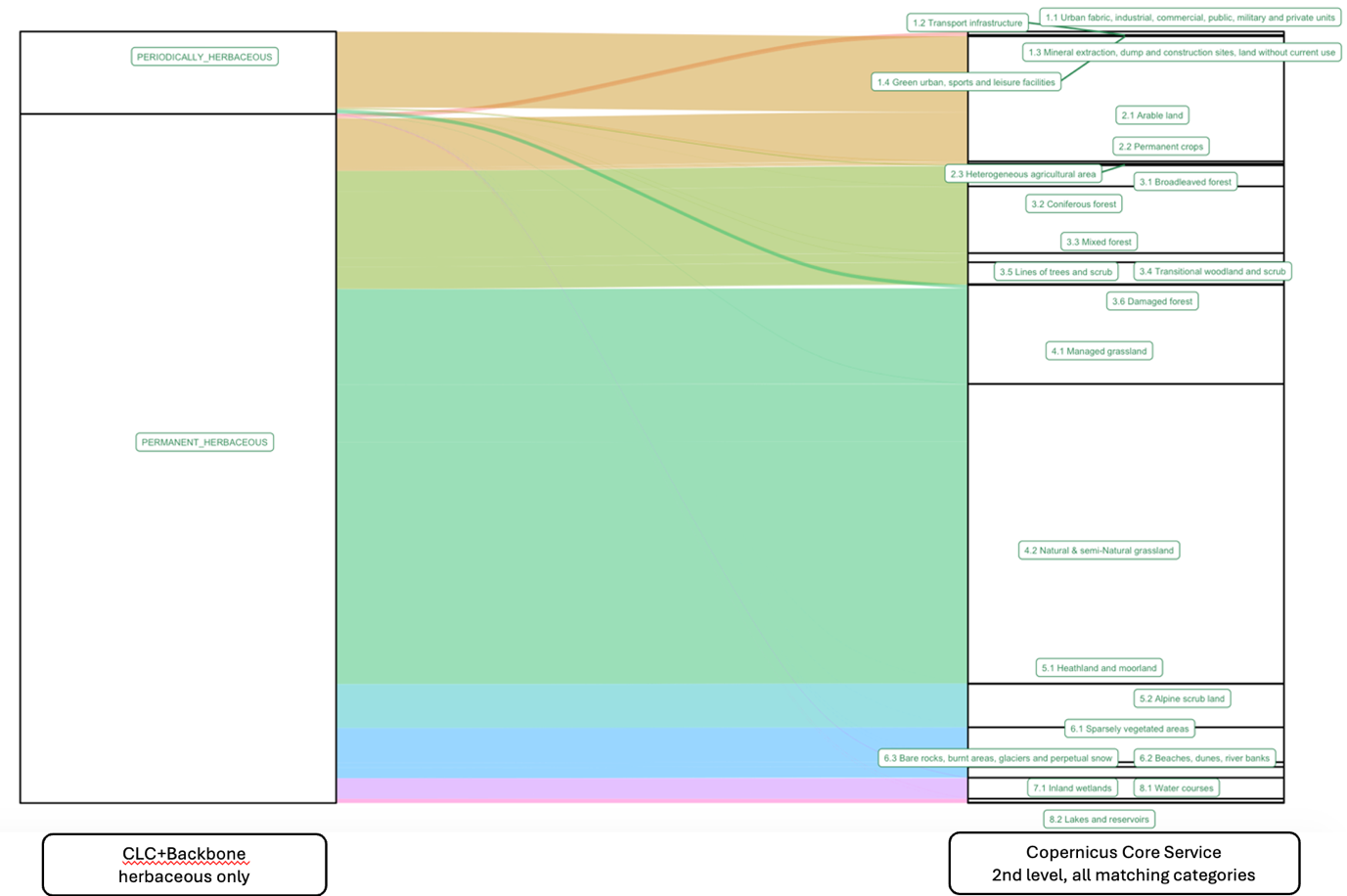

**Figure S6.4:** Testing which polygon categories are found in the CLC categories permanent and periodically herbaceous confirms that the periodically herbaceous category consists almost exclusively of arable land and greenhouses, while the permanent herbaceous class is a mix of arable land, managed and natural grassland, shrubland, and different forest types.

Because sealed surfaces cover only a small area, we created close-ups of all flows from the urban classification at the first level of the polygon data to all raster classifications (Figure S6.5) and all flows from the sealed classification in the raster data to all polygon classifications (Figure 2.6.6). The flow diagrams show that many urban classes in the polygon data include flows to natural classifications in the raster data. Flows from transport infrastructure to different types of trees exist, as well as flows from the urban fabric to permanent herbaceous, likely representing parks and gardens. At the same time, flows from the sealed classification in the raster data can be found in several second-level classifications of all 8 main classes in the polygon data, with the built-up classification contributing about 50%. However, even though mismatches in the sealed category of the satellite imagery are likely to exist, the satellite data seems better suited to make the distinction between built-up and green spaces, as the urban category in the polygon data shows strong flows to both sealed and permanent herbaceous in the raster layer (Figure S6.5).
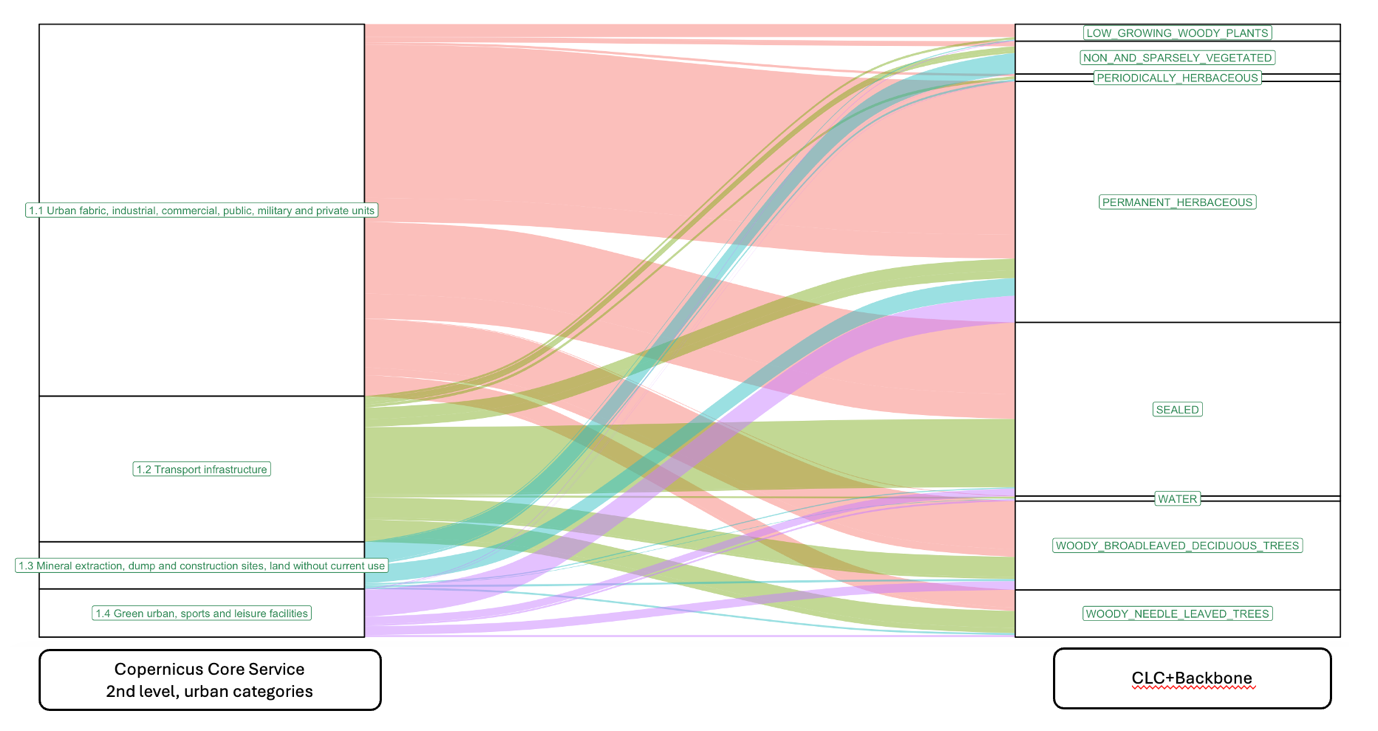

**Figure S6.5:** The satellite imagery (CLC+) seems more likely to capture sealed surfaces than the polygon category “urban” at levels 1 and 2. The urban/build-up class in the polygon dataset classifies military areas and parks together with urban areas. In the CLC raster, military areas and parks likely get classified as the natural vegetation type that is found in these locations. Similarly, urban fabric in the polygon data likely includes parks and gardens based on the strong flow to the permanent herbaceous category in the raster data.

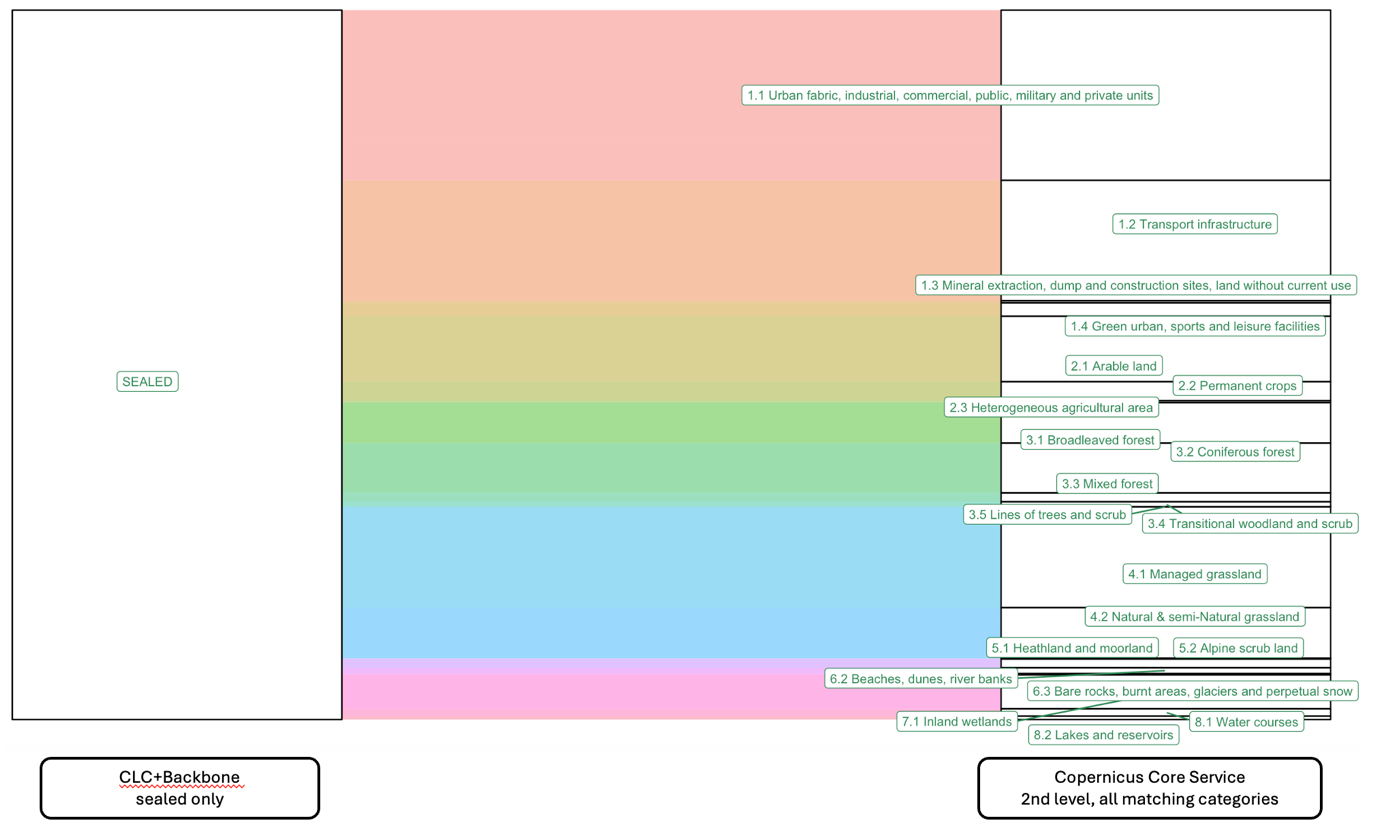

**Figure S6.6:** Flows from the raster category “Sealed” to all polygon classifications. The built-up and agricultural sub-classifications (starting with 1. and 2.) make up about 50% of the flows, and pixels classified as “sealed” in all other main polygon classifications in varying quantities.
